# USP7 maintains hematopoietic stem cell dormancy and function by stabilizing HMGA2

**DOI:** 10.64898/2026.07.03.731117

**Authors:** Antoine Nouhaud, Anna Diaz, Mathieu Bouttier, Quentin Rigaud, Pauline Enfedaque, Hedi Somai, Sylvie Hebrard, Naïs Prade, Stéphanie Dufrechou, Daniele Musiani, Mariette Matondo, Guillaume Andrieu, Marlène Pasquet, Laetitia Largeaud, Cyril Broccardo, Éric Delabesse, Bastien Gerby, Christine Didier

## Abstract

Hematopoietic stem cell (HSC) longevity critically depends on maintaining a deep dormant state, yet the molecular mechanisms that preserve this rare and functionally essential population remain poorly understood. Here, we identify the deubiquitinase USP7 as a key regulator of long-term HSC dormancy.

Using a *Usp*7^+/-^ mouse model, we uncover selective depletion of hematopoietic stem and progenitor cells (HSPCs), which is associated with impaired long-term repopulation capacity. Strikingly, H2B-GFP label-retention assays reveal a profound loss of dormant HSCs in *Usp*7^+/-^ mice, demonstrating a failure to maintain the most quiescent stem cell fraction *in vivo*. Consistently, single-cell RNA sequencing shows erosion of the transcriptional dormancy program, linking USP7 activity to the preservation of stem cell identity at both functional and molecular levels. Mechanistically, ultra-low-input proteomic profiling and biochemical approaches identify HMGA2 as a novel USP7 substrate, suggesting that ubiquitin-dependent regulation of chromatin architecture contributes to the control of HSC dormancy.

Together, our findings establish USP7 as a critical regulator of HSC dormancy, revealing a previously unrecognized post-translational mechanism controlling stem cell longevity, with implications for aging, regeneration, and hematopoietic disorders.

## INTRODUCTION

Hematopoietic stem cells (HSCs) possess the unique capacity for self-renewal and multilineage differentiation, enabling the lifelong generation of both daughter stem cells and committed progenitors, thereby sustaining hematopoietic homeostasis *(de Haan et Lazare 2018; Sean J. Morrison et Spradling 2008)*. This delicate balance between quiescence, self-renewal, and differentiation is tightly regulated, and its disruption can lead to stem cell exhaustion or malignant transformation, ultimately giving rise to leukemia-initiating stem cells (LSCs) *(Flach et al. 2014)*. Understanding the mechanisms that preserve HSC homeostasis is therefore fundamental both for deciphering the regulation of normal hematopoiesis and for developing strategies to selectively target LSCs.

Under steady-state conditions, HSCs reside predominantly in a quiescent state within specialized bone marrow niches and can only be activated by external stimuli to exit quiescence and enter the cell cycle *(Trumpp et al. 2010)*. Historically, HSCs have been classified into two main functional subtypes: long-term HSCs (LT-HSCs), which sustain lifelong hematopoiesis, and short-term HSCs (ST-HSCs), which possess more limited self-renewal potential *(S. J. Morrison et Weissman 1994; Seita et Weissman 2010)*. However, seminal studies from the Trumpp laboratory revealed that the LT-HSC compartment itself is heterogeneous. Comprising both dormant HSCs (dHSCs) and active HSCs (aHSCs), and notably the dHSCs displaying superior serial engraftment capacity in transplantation assays *(A. Wilson et al. 2008; Foudi et al. 2009; Qiu et al. 2014)*. Dormant HSCs reside in a deeply quiescent state characterized by transcriptional and metabolic repression *(Cabezas-Wallscheid et al. 2017; Bartram et al. 2026).* These cells are arrested in the G_0_ phase of the cell cycle and exhibit extremely low biosynthetic activity *(Cabezas-Wallscheid et al. 2014; Rettkowski et Cabezas-Wallscheid 2025)*. Experimentally, dHSCs can be identified using label-retention approaches or reporter mouse models such as H2B-GFP or Gprc5c-GFP*(A. Wilson et al. 2008; Cabezas-Wallscheid et al. 2017)*. Maintenance of this deep quiescence state is critical to preserve stem cell integrity, prevent stem cell exhaustion, and ensure appropriate hematopoietic lineage output throughout life.

Post-translational regulation is emerging as a central mechanism that fine-tunes the balance between HSC self-renewal and differentiation *(Signer et al. 2014; Lv et al. 2021)*. The ubiquitin-proteasome system, through ubiquitination and deubiquitination, dynamically controls protein stability, localization, and activity, thereby shaping key cellular programs. Ubiquitin-specific proteases (USPs) are important regulators of HSC maintenance, in part by stabilizing proteins that govern cell cycle progression, DNA replication, and chromatin organization (USP1, USP3, USP4, USP7, USP10, USP15, USP16 and CYLD *(Parmar et al. 2010; Lancini et al. 2014; Liu et al. 2024; Shan et al. 2026; Higuchi et al. 2016; van den Berk et al. 2020; Gu et al. 2016; Tesio et al. 2015)*. Dysregulation of these pathways compromises stem cell fitness and hematopoietic homeostasis and can predispose to malignant transformation.

Among USPs, USP7 has emerged as a pleiotropic regulator with broad oncogenic relevance. USP7 stabilizes multiple substrates, including the E3 ligase MDM2 and its target p53 *(Li et al. 2004)*, DNA damage response factors such as Claspin *(Faustrup et al. 2009)*, and chromatin regulators including DNMT1*(Qin et al. 2011)* and the histone demethylase PHF8 *(Q. Wang et al. 2016)*. Through these activities, USP7 integrates stress signaling, transcriptional regulation, and epigenetic control to influence cell proliferation, differentiation, and survival. In leukemia, this regulatory axis is frequently hijacked to sustain aberrant self-renewal and block differentiation. Elevated *USP7* expression or activity has been reported in both acute myeloid leukemia (AML) and acute lymphoblastic leukemia (ALL), where it promotes leukemic cell survival, maintains oncogenic transcriptional programs, and contributes to chemotherapy resistance *(Cartel et al. 2021; Shan et al. 2018)*. Despite its emerging relevance in hematologic malignancies, the role of USP7 in normal HSC biology remains understudied, particularly regarding the regulation of quiescence and long-term stem cell maintenance. In this study, we show that USP7 functions as a deubiquitinase targeting HMGA2, revealing a previously unknown mechanism that preserves HSC quiescence and stem cell function. Our findings provide fundamental insights into the post-translational control of hematopoietic stem cell homeostasis and highlight potential vulnerabilities that may be exploited in leukemia.

## RESULTS

### USP7 maintains hematopoietic stem and progenitor cells with limited effects on mature blood compartments

We first assessed USP7 expression across normal mouse hematopoiesis using publicly available transcriptomic and proteomic datasets; (Figure S1A-B). USP7 is the third most abundant USP expressed in the stem cell compartment (Figure S1A) and is expressed homogeneously across the hematopoietic hierarchy, with comparable levels in LT-HSCs and early lineage-committed progenitors (Figure S1B). Notably, *Usp7* expression is markedly upregulated during terminal erythroid differentiation. This expression pattern is conserved in humans, as *USP7* is similarly expressed in CD34⁺ HSCs, early progenitors and across the whole human hematopoietic hierarchy (Figure S1C).

We then generated constitutive heterozygous *Usp7* knockout (*Usp7*^+/-^) mice, to model physiological gene dosage and clinically relevant loss-of-function alterations. These mice displayed no visible detrimental phenotype and showed bone marrow (BM) cellularity comparable to wild-type controls (Figure S1D). As expected, USP7 protein in the BM was reduced by ∼40% compared with wild-type controls (Figure S1E), with a corresponding decrease in RNA expression by ∼36% in *Usp7*^+/-^ mice (Figure S1F). To comprehensively characterize hematopoietic populations from stem cells to mature lineages, we developed a multiparametric spectral flow cytometry panel incorporating surface markers spanning the full spectrum of hematopoietic stem and progenitor cells (HSPCs) and differentiated cells (Figure 1A; S1G) and Table S1A. We further established an unsupervised clustering framework based on Uniform Manifold Approximation and Projection (UMAP) to reconstruct the hematopoietic landscape, enabling comprehensive visualization of differentiation across all lineages, including a dedicated embedding for the HSPC compartment. In both representations, populations are color-coded according to phenotypic identities defined by gating strategies (Figure 1B). A schematic overview of the HSPC compartment is provided (Figure 1C). Overlay of marker expression onto the HSPC UMAP supported cluster annotation (Figure 1D). As expected, CD127 expression was restricted to the common lymphoid progenitors (CLPs) compartment, whereas CD105 expression was enriched in the long-term hematopoietic stem cells (LT-HSCs) and megakaryocytic-erythroid progenitors (MEPs), consistent with established phenotypic definitions *(Meurer et Weiskirchen 2020)*. Application of spectral cytometry showed that mature hematopoietic compartments were largely preserved, with the exception of CD8⁺ T lymphocytes and megakaryocytes (MKs), which were selectively reduced in the bone marrow of *Usp*7^+/-^ mice compared with wild-type controls (Figure S1H). Detailed analysis of B-cell development highlighted no major alterations except for the Pro-B fraction (Figure S1I), indicating that steady-state hematopoietic differentiation remains largely intact upon *Usp7* haploinsufficiency. Peripheral blood analysis further showed a significant decrease in total white blood cells and lymphocytes in *Usp*7^+/-^ mice (Figure S1J). Deep phenotypic profiling revealed a significant reduction in Lin⁻ and Lin⁻Kit^+^Sca⁻ (LK) progenitor populations in the bone marrow of 3-5-month-old *Usp*7^+/-^ mice (Figure 1B (right panel); 1E). Notably, the absolute numbers of all committed progenitors, including common myeloid progenitors (CMPs), granulocyte-macrophage progenitors (GMPs), megakaryocytic-erythroid progenitors (MEPs), and common lymphoid progenitors (CLPs) were reduced (Figure 1F). This reduction extended to the Lin^−^Sca-1^+^c-Kit^+^ (LSK) compartment (Figure 1E), which was markedly diminished in *Usp*7^+/-^ mice, with a pronounced decrease in multipotent progenitor subsets, including MPP6, MPP5, MPP3, and MPP4 (Figure 1G). At the apex of the hematopoietic hierarchy, the LT-HSC compartment remained unaltered in *Usp*7^+/-^ mice.

**Figure 1.**
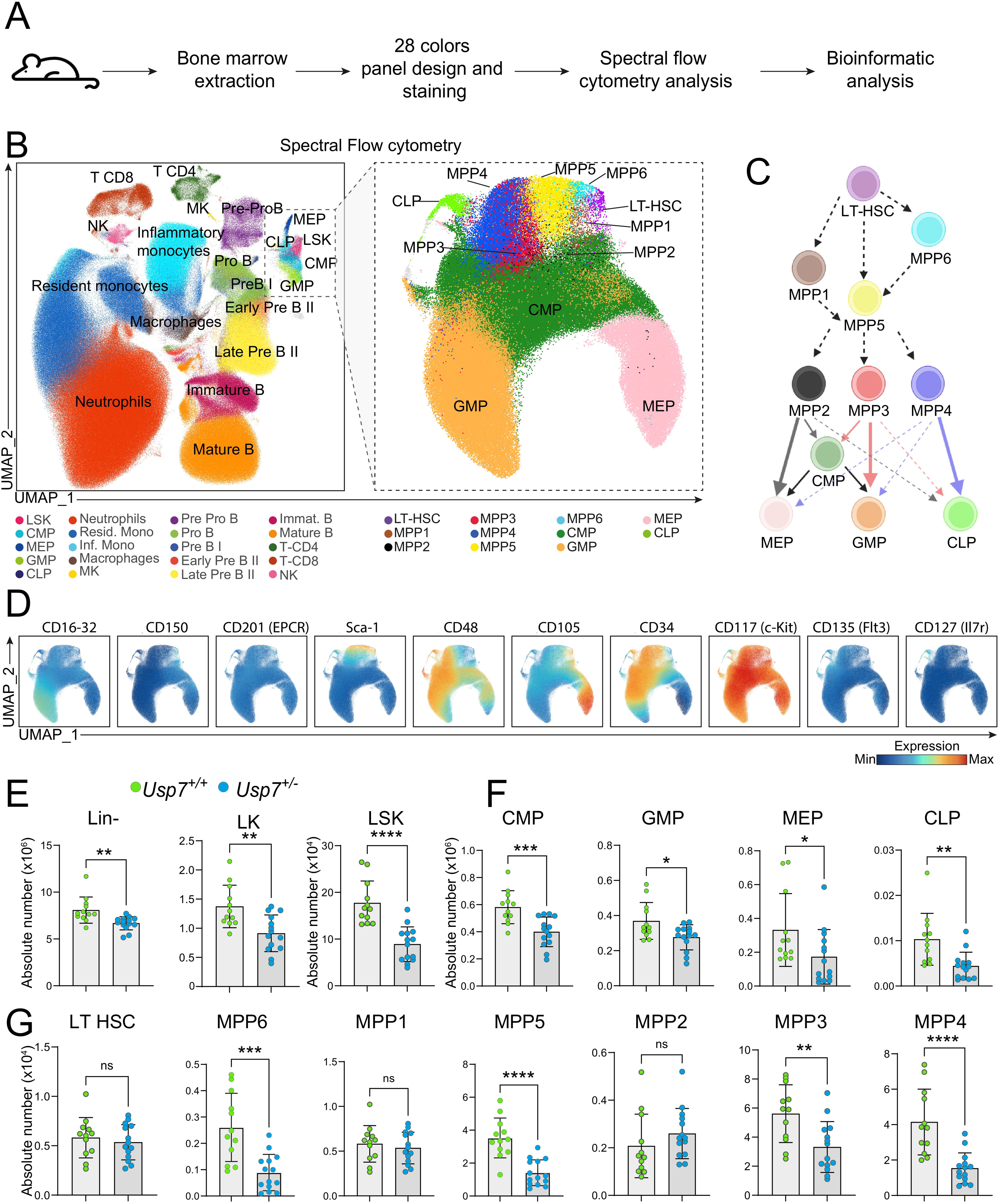
USP7 maintains hematopoietic stem and progenitor cells with minor effects on mature compartments. (**A**) Experimental workflow of phenotypic characterization of the hematopoietic compartments from *Usp7*^+/+^ and *Usp7*^+/-^ mice by spectral cytometry (**B**) UMAP embedding of spectral flow cytometry data, with cells colored according to gating defined identities (gating strategy shown in Figure S1G). *Left panel*, global view of non-erythroid bone marrow hematopoietic cells. *Right panel*, hematopoietic stem and progenitors cell compartments. (**C**) Hierarchical organization of hematopoietic progenitors. Schematic representation of differentiation trajectories from LT-HSCs through multipotent progenitors (MPP1-6) toward lineage-committed progenitors, including common myeloid progenitors (CMPs), granulocyte-monocyte progenitors (GMPs), megakaryocyte-erythroid progenitors (MEPs), and common lymphoid progenitors (CLPs). Arrows indicate inferred lineage relationships (created with BioRender.com). (**D**) UMAP representation of hematopoietic stem and progenitor cells (HSPCs) colored according to the expression level of CD16-32, CD150, CD201 (EPCR), Sca-1 CD48, CD105, CD34, CD117(c-Kit), CD135 (Flt3) and CD127 (Il7r). (**E-G**) Absolute numbers of hematopoietic stem and progenitor cell populations in 3-5-month-old *Usp7*^+/+^ (green) and *Usp7*^+/-^ (blue) mice. (**E**) Lin⁻, LK (Lin⁻ Sca1⁻ c-Kit⁺), and LSK (Lin⁻ Sca1⁺ c-Kit⁺) compartments (**F**) CMPs, GMPs, MEPs, and CLPs. (**G**) Long-term hematopoietic stem cells (LT-HSCs) and multipotent progenitors’ subsets (MPP1-6). Data are shown for two independent experiments (n = 12 *Usp7*^+/+^ and n = 14 *Usp7*^+/-^ mice). Results are shown as median ± SD; ns not significant, *p < 0.05, **p < 0.01, ***p < 0.001, ****p < 0.0001.

Altogether, these data demonstrate that USP7 plays a critical role regulating the number of hematopoietic stem and progenitor cells.

### USP7 sustains HSPCs engraftment potential

To explore the role of USP7 in HSPCs at the functional level, we first performed serial competitive transplantation assays. Total BM from *Usp7*^+/-^ donor mice or control littermates (CD45.2) was transplanted into CD45.1 recipient mice, and engraftment efficiency was quantified several months after primary and secondary transplantation (Figure S2A-B). We found that *Usp7*^+/-^ BM cells exhibited reduced repopulation capacity compared to control counterparts in primary and secondary transplantations (Figure S2B-C). Following primary transplantation, the contribution of T cells, B cells, and myeloid lineages to total donor-derived cells was comparable between groups. In contrast, upon secondary transplantation, a progressive skewing toward myeloid differentiation, accompanied by reduced T- and B-cell output, was observed in *Usp7*^+/-^ chimeras over time (Figure S2D). These results indicate that USP7 is required to maintain balanced hematopoietic differentiation under regenerative stress.

To further investigate the impact of USP7 on repopulation capacity, we performed competitive transplantation using *Usp7*⁺^/+^ or *Usp7*^+/-^ LSK from CD45.2 mice, LSK cells from CD45.1 mice combined with support CD45.1/2 BM cells into lethally irradiated CD45.1/2 recipients (Figure 2A). Primary recipients transplanted with *Usp7*^+/-^ LSK exhibited decreased chimerism from 1 to 4 months post-transplantation compared to controls (Figure 2B-C). Secondary transplantation of total bone marrow from the primary transplanted mice confirmed the loss of long-term engraftment capacity of *Usp7***^+/-^**cells (Figure 2B-C).

**Figure 2.**
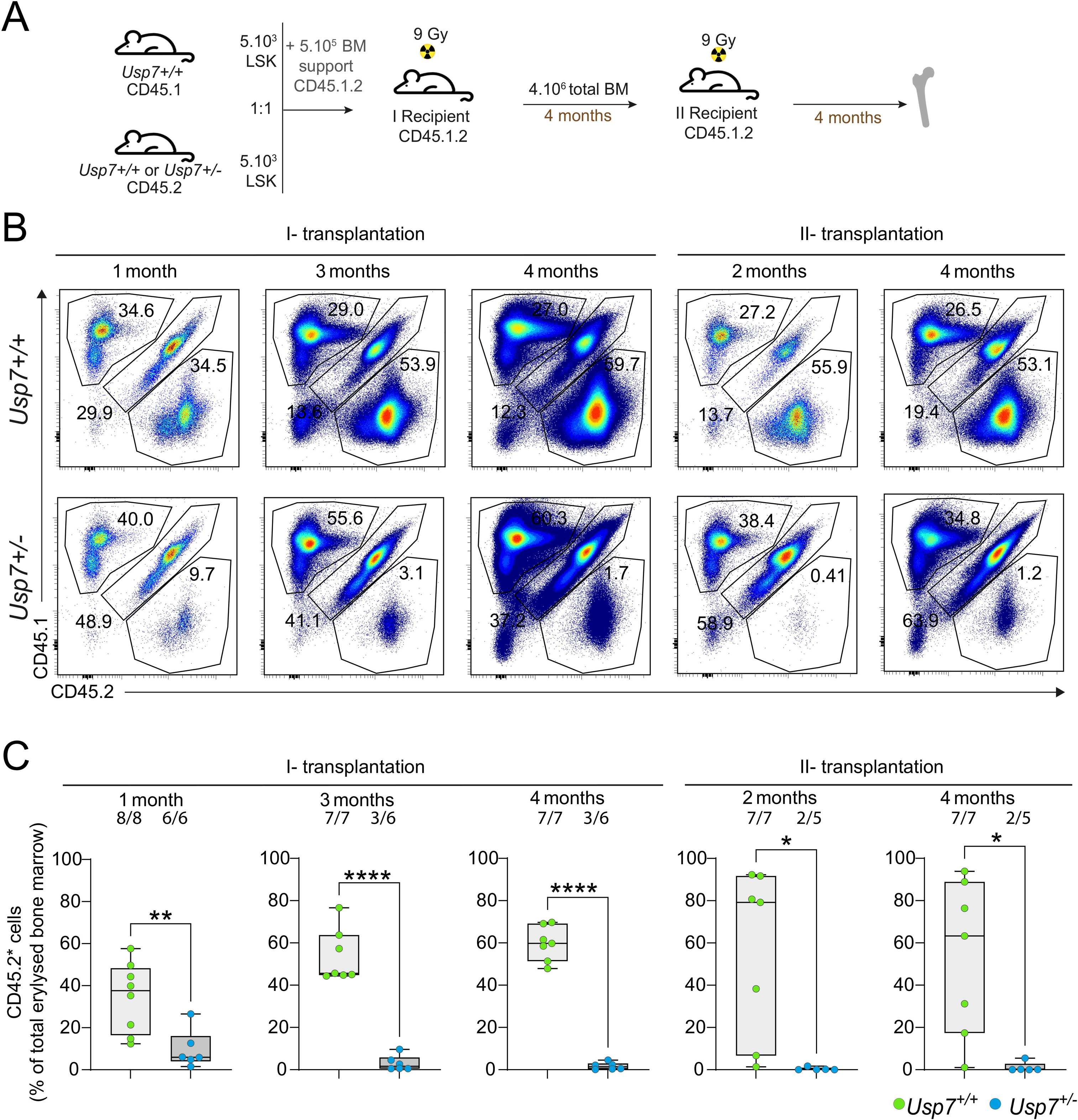
USP7 regulates engraftment potential of HSPCs. (**A**) Experimental workflow of competitive transplantation assays. LSK cells from 2-month-old CD45.1 wild-type mice were mixed with LSK cells from 2-month-old CD45.2 *Usp7*^+/+^or *Usp7*^+/-^ mice, together with CD45.1/2 BM support cells, and transplanted into 3-4-month-old CD45.1/2 recipient mice (n=8 recipients for *Usp7*^+/+^ : *Usp7*^+/+^competition; n=6 recipients for *Usp7*^+/-^ : *Usp7*^+/+^ competition). For secondary transplantation, total BM cells from primary recipients were injected into new 3-month-old CD45.1/2 mice recipient mice (n=7 recipients for *Usp7^+/+^* : *Usp7^+/+^*competition; n=5 recipients for *Usp7*^+/-^ : *Usp7*^+/+^ competition). (**B**) Representative flow cytometry density plots of concatenated bone marrow samples, gated on Ter119**⁻** cells, showing donor-derived engraftment at 1, 3 and 4 months after the first transplantation, and at 2 and 4 months after second transplantation. (**C**) Quantification of donor CD45.2⁺ engraftment at 1, 3 and 4 months after primary transplantation (left side of the panel), and at 2 and 4 months after secondary transplantation (right side of the panel). The number of engrafted mice (> 1% of CD45.2⁺ cells) out of the total transplanted mice are indicated above the graph. Each green dot represents an individual *Usp7*^+/+^mouse, and each blue dot represents a *Usp7*^+/-^ mouse. Results are shown as median ± SD; ns not significant, *p < 0.05, **p < 0.01, ***p < 0.001, ****p < 0.0001.

Collectively, these results suggest that *Usp7*^+/-^ HSPCs display impaired long-term repopulation capacity.

### Single-Cell Molecular Profiling Reveals Altered Programs in *Usp7*^+/-^ HSCs

To investigate the molecular basis underlying the impaired long-term repopulation capacity caused by *Usp7* haploinsufficiency, we performed single cell RNA sequencing (scRNA-seq) of LSK cells from *Usp7*^+/-^ and wild-type mice. Unsupervised clustering using UMAP identified transcriptionally distinct clusters within the LSK compartment, ranging from hematopoietic stem cells (HSCs) to more primed progenitors (Figure 3A). Because cell-cycle-associated transcription strongly influenced clustering, we regressed out cell-cycle effects prior to downstream analyses to ensure that clusters reflected true lineage-priming states rather than proliferation status (Figure S3A).

**Figure 3.**
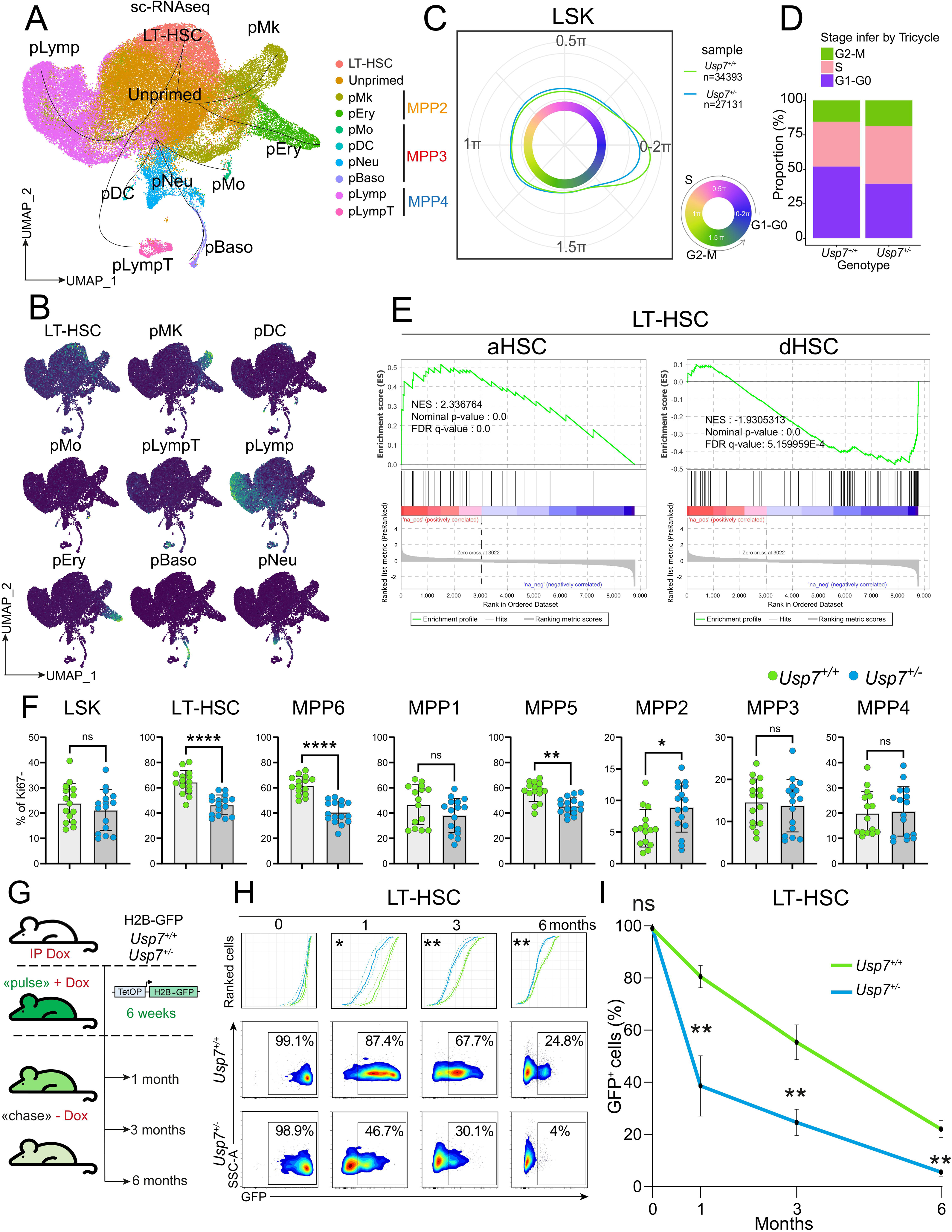
USP7 is essential for maintaining HSC dormancy. (**A**) UMAP representation of single-cell RNA-seq data from LSK cells from *Usp7*^+/+^ and *Usp7*^+/-^ mice after cell cycle regression and cluster annotation. Colors correspond to groups identified through annotated clusters. (**B**) UMAP plots of LSK cells colored by the expression of lineage-specific gene signatures and reference datasets (Table S3), illustrating the localization and distribution of these markers across different LSK differentiation (e.g., LT-HSC, pMK, pDC). (**C**) Representation of cell cycle status of LSK cells from *Usp7*^+/+^ (n=34,393) and *Usp7^+/-^*(n=27,131) using the Tricycle package *(S. C. Zheng et al. 2022)*. Estimated cell cycle positions were bounded between 0 and 2π. Phases were mapped onto a circular scale (θ, radians): S phase at 0.1π-π, G2/M at π-1.75 π, and G1/G0 spanning 1.75π-0.1π, providing a continuous representation of cell cycle progression. (**D**) Percentage of LSK cells in each cell cycle phase. G0-G1 (purple), S (pink), and G2M (green). (**E**) GSEA analysis of activated HSC (aHSC) and dormant HSC (dHSC) *(Cabezas-Wallscheid et al. 2017)* in LT-HSC cluster from *Usp7*^+/+^ and *Usp7*^+/-^ mice. Positive NES indicates enrichment in *Usp7*^+/-^ LT-HSCs, whereas negative NES indicates enrichment in *Usp7*^+/+^ LT-HSCs. (**F**) Percentage of Ki-67⁻ (G0) cells in LSKs, LT-HSCs, and MPP1-6 subsets from *Usp7*^+/+^ (n=15) and *Usp7*^+/-^ (n=16) mice. 3 independent experiments. (**G**) Experimental procedure to assess *in vivo* cell division kinetics in *Usp7*^+/+^ and *Usp7*^+/-^ mice (n = at least 3). Mice were sacrificed and analyzed at 0, 1, 3, and 6 months of DOX chase. (**H**) Representative FACS analysis of GFP expression in the LT-HSCs from *Usp7*^+/+^ (central panels) and *Usp7*^+/-^ (bottom panels) mice for each time point of DOX chase. AUC (area under the curve) quantification of the GFP (upper panels) was compared between the two conditions. (**p < 0.01, ***p < 0.001). (**I**) The proportion of GFP-positive cells is indicated for each time point. Results are shown as median ± SD; ns not significant, **p < 0.01.

We then identified and annotated these clusters by integrating published gene expression signatures *(Rodriguez-Fraticelli et al. 2018)* (Table S3) (Figure 3A–B; S3B), which allowed us to merge transcriptionally related clusters into defined cell-state populations. Gene set scoring of lineage-specific expression signatures, derived from published datasets, revealed robust and specific enrichment of each signature within its corresponding annotated cluster, with maximal average signature scores observed in the expected cell populations, including LT-HSC-associated signatures peaking in the LT-HSC cluster (Figure S3C). While the global topology of the UMAP was preserved between genotypes (Figure S3B), comparative density analysis (Figure S3D) showed a redistribution of cell states. Specifically, the *Usp7*^+/-^ LSKs exhibited a decrease in the lymphoid-primed cluster compared to their wild-type counterparts. Comparison of transcriptional clusters with CaSTLe-based *(Lieberman et al. 2018)* cell identity assignments showed a high level of concordance using published inDrops reference datasets *(Rodriguez-Fraticelli et al. 2018)*, with approximately 78% of cells in the LT-HSC cluster classified as LT-HSCs by the algorithm (Figure S3E-G).

We further analyzed cell-cycle dynamics in each genotype using the tricycle package *(S. C. Zheng et al. 2022)*. Circular embedding onto the transcriptional cell-cycle manifold revealed that *Usp7*^+/-^ LSK cells preferentially accumulated in S and G2/M phases, with reduced G0/G1 distribution compared to wild-type cells (Figure 3C-D). Examination of canonical cell-cycle genes confirmed these positions: *Pcna*, for example, peaked in S phase (Figure S3H), consistent with its known expression profile. Together, these data indicate that *Usp7* haploinsufficiency promotes cell cycle progression.

Furthermore, differential gene expression analysis (DEGs) revealed that *Usp7*^+/-^ LT-HSCs preferentially upregulated genes linked to cell cycle progression (*Hist1h2ap*) and downregulated genes associated with stemness (*Meg3, Meis1, Malat1*). Notably, *Nupr1* expression was significantly increased in *Usp7*^+/-^ LT-HSCs, consistent with prior studies demonstrating that *Nupr1* deficiency enhances HSC engraftment efficiency and functional output *(T. Wang et al. 2022)* (Figure S3I and Table S4). This transcriptional shift mirrors the results of Gene Set Enrichment Analysis (GSEA) performed on the DEGs, in which the active HSC (aHSC) signature was enriched in *Usp7*^+/-^ HSCs, whereas the dormant HSC (dHSC) signature *(Cabezas-Wallscheid et al. 2017)* was enriched in wild-type cells (Figure 3E; Table S3, S4 and S8).

Overall, these single-cell analyses reveal that *Usp7* haploinsufficiency shifts HSCs toward a more proliferative and primed transcriptional state, at the expense of dormancy and stemness programs.

### USP7 controls LT-HSC dormancy and restrains cell-cycle entry

Building on these transcriptomic insights, we assessed Ki-67 expression in LSK compartments from *Usp7*^+/+^ and *Usp7*^+/-^ mice under steady-state conditions (Figure 3F; S3J). LT-HSCs, MPP6 and MPP5 from *Usp7*^+/-^mice exhibited significantly fewer quiescent (G0) cells compared to wild-type cells.

To further address the impact of USP7 on LT-HSC dormancy, we generated *Usp7*^+/-^ ::*H2B-GFP*^tg^ mice, by crossing *Usp7* heterozygous animals with a doxycycline-inducible *H2B-GFP*^tg^ mouse model, a well-established approach to monitor stem cell quiescence *in vivo (Foudi et al. 2009; A. Wilson et al. 2008)*. Following a doxycycline pulse, cell division kinetics were monitored *in vivo* in each compartment by tracking the progressive dilution of GFP fluorescence over time (Figure 3G-I) using a multiparametric spectral flow cytometry staining strategy (Figure S3K-L; Table S1E). At the end of the pulse, nearly all hematopoietic cells in both *Usp7*^+/-^ and wild-type mice expressed H2B-GFP, confirming efficient and comparable labeling between genotypes (Figure 3H; S3L). As expected, under wild-type conditions, LT-HSCs displayed the highest GFP retention, followed closely by MPP6, and then by MPP1 and MPP5. In contrast, other MPP populations (MPP2-4) exhibited markedly lower GFP retention, while committed progenitors rapidly lost GFP signal within one month after doxycycline (DOX) withdrawal (Figure 3H; S3L). Strikingly, GFP fluorescence declined more rapidly in LT-HSCs from *Usp7*^+/-^ mice compared with wild-type controls as well as in MPP1, MPP5, and MPP6, indicating a higher proliferative rate in *Usp7*-deficient cells (Figure 3H-I; S3L). Consistent with previous reports that dormant HSCs (dHSCs) divide approximately once every 145 days *(A. Wilson et al. 2008)*, HSCs retaining high H2B-GFP levels six months after labeling were defined as dHSCs. These GFP-positive LT-HSCs also displayed more EPCR expression than GFP-negative cells, which is known to be preferentially expressed in quiescent LT-HSCs *(Rabe et al. 2020; Lin et Trumpp 2023)*, in both genotypes and at each time point after chase (Figure S3M-N).

Altogether, these data establish USP7 as critical for preserving LT-HSC dormancy, with *Usp7*^+/-^ HSCs showing both accelerated division and impaired stem cell identity, paralleling functional results.

### USP7 preserves LT-HSCs function by modulating molecular pathways and regeneration capacity

To further assess the function of *Usp7*-deficient LT-HSCs, we performed RNA-sequencing (RNA-seq), proteomics, and functional transplantation assays (Figure 4A). LT-HSCs were isolated from *Usp7*^+/+^ and *Usp7*^+/-^ mice using the gating strategy (Figure 4B; S4A). We transplanted LT-HSCs in competition alongside CD45.1/2 wild-type BM cells into lethally irradiated CD45.1 recipients (Figure 4C). Strikingly, *Usp7*^+/-^ LT-HSCs showed almost no detectable engraftment, markedly more impaired than total LSK cells. Analysis of primary chimeras at 1 and 7 months post-transplant confirmed a complete loss of reconstitution capacity, demonstrating that *Usp7* haploinsufficiency abolishes LT-HSC self-renewal.

**Figure 4.**
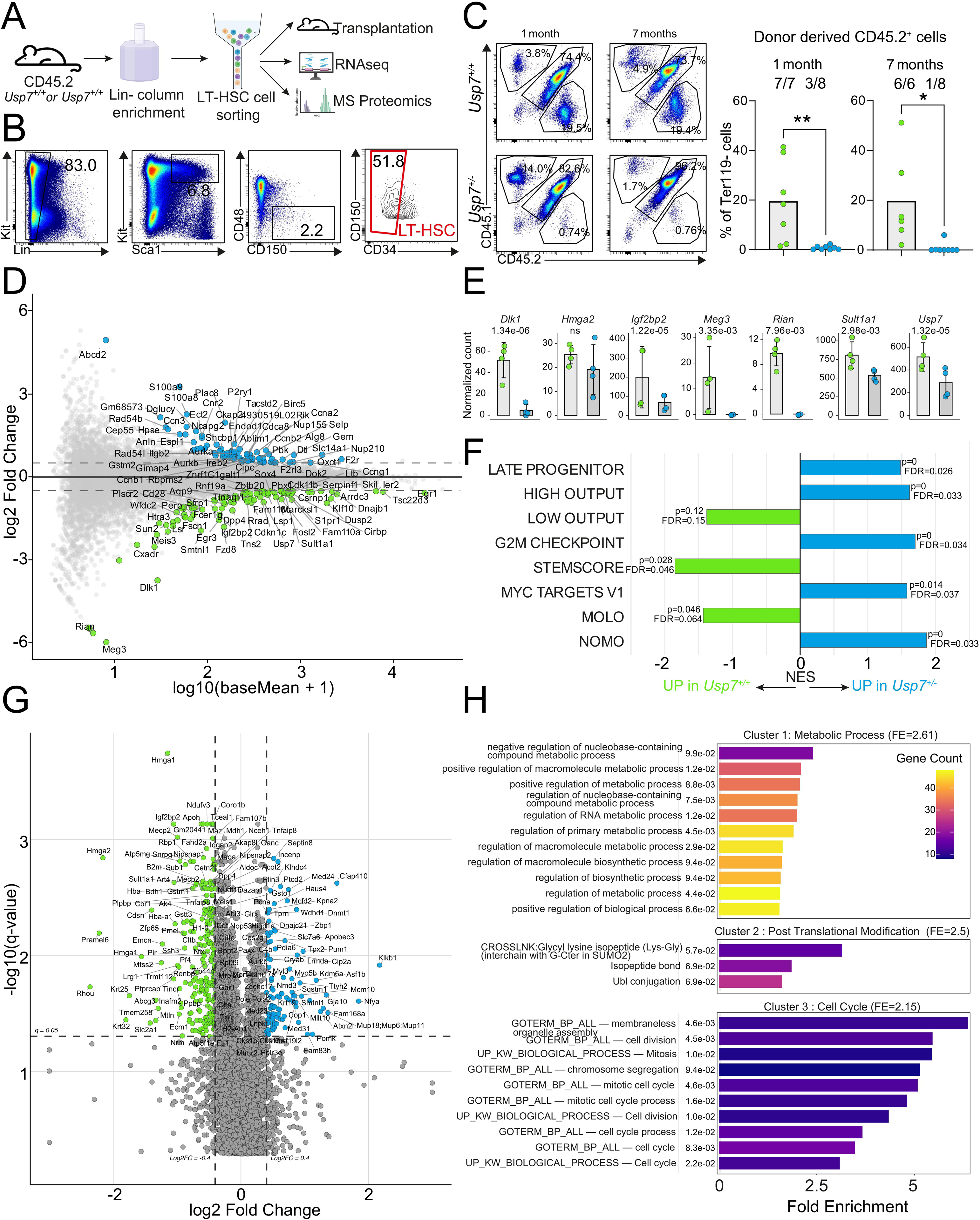
USP7 preserves LT-HSC function and quiescence molecular programs. (**A**) Workflow of functional and molecular analysis of LT-HSCs (created with BioRender.com). (**B**) Gating strategy to sort LT-HSCs. (**C**) A total of 250 LT-HSCs from *Usp7*^+/+^ and *Usp7*^+/-^ CD45.2 donor mice were mixed with 1.10^6^ CD45.1/2 bone marrow support cells and transplanted into CD45.1 recipients. Representative flow cytometry density plots of concatenated BM samples gated on Ter119⁻ live singlets, showing CD45.2⁺ donor-derived hematopoietic engraftment at 1 and 7 months post-transplantation (left panel). Quantification of CD45.2⁺ donor chimerism in the BM at both time points is presented (right panel). Data represent mean ± SD (n = 6-8 mice per group). Statistical analyses were performed using a two-tailed t-test; **p < 0.01, ***p < 0.001. (**D**) MA-plot depicting differential gene expression in *Usp7*^+/+^ versus *Usp7*^+/-^ LT-HSCs from ultra-low input RNA-sequencing (*n*= 4 independent biological replicates per genotype). Each point represents a gene, with the x-axis showing expression strength (log10(baseMean + 1)) and the y-axis showing the log2FC. Genes with p-val-adj < 0.05 and |log2FC| > 0.5 are highlighted: upregulated in *Usp7*^+/-^ (blue) and downregulated (green); non-significant genes are shown in light grey. Dashed horizontal lines mark the log2FC thresholds (±0.5). (**E**) Bar-plot represents mean normalized counts ± SD, with individual samples shown as points in *Usp7^+/+^ versus Usp7^+/-^* LT-HSCs. P-val-adj are indicated. (**F**) Gene Set Enrichment Analysis (GSEA) of literature-derived signatures (see Table S3) comparing *Usp7*^+/+^ and *Usp7*^+/-^ LT-HSCs highlights differential expression of key pathways. Positive NES indicates enrichment in *Usp7*^+/-^ LT-HSCs, whereas negative NES indicates enrichment in *Usp7*^+/+^ LT-HSCs (**G**) Volcano-plot showing differential protein expression in *Usp7*^+/+^ *versus Usp7*^+/-^ LT-HSCs. Log₂FC are plotted against q-value (q) from ultra-low input MS/MS analysis (*n* = 4). Proteins significantly upregulated in *Usp7*^+/-^ LT-HSCs (log₂FC > 0.4, q < 0.05) are highlighted in blue, whereas significantly downregulated proteins (log₂FC < −0.4, q < 0.05) are shown in green. (**H**) DAVID analysis on proteins significantly upregulated in *Usp7*^+/-^ *versus Usp7*^+/+^ LT-HSCs (log₂FC > 0.3; q < 0.05). Enriched pathways are shown with Fold Enrichment (FE), false discovery rate (FDR) and p(Benjamini).

To explore the molecular basis of the defect, we performed RNA-seq analysis of *Usp7*^+/+^ and *Usp7*^+/-^ LT-HSCs. Principal component analysis (PCA) of the transcriptome profiles revealed a clear segregation between wild-type (WT) and heterozygous (HT) LT-HSCs, highlighting genotype-dependent transcriptional remodeling (Figure S4B). Differential expression analysis (Table S7) revealed a distinct set of genes significantly up- or downregulated in *Usp7* ^+/-^ LT-HSCs compared to wild-type controls (Figure 4D). As expected, *Usp7* expression was reduced by approximately 50% in *Usp7**^+/-^*** LT-HSCs compared to wild-type samples. Furthermore, *Dlk1*, *Igf2bp2*, *Meg3*, *Rian* and *Sult1a1* genes known to correlate with HSC self-renewal capacity or quiescence *(Cabezas-Wallscheid et al. 2014, 2017; Sommerkamp et al. 2019; Huang et al. 2023; Suo et al. 2022; Kubota et al. 2024)* were found to be significantly downregulated in *Usp7***^+/-^**LT-HSCs compared to wild-type samples (Figure 4E). GSEA revealed significant enrichment of cell-cycle-related gene sets, including Hallmark G2/M Checkpoint and Myc Targets V1 *(Howe et al. 2018)*. It also showed enrichment of signatures associated with activated or differentiated HSCs, such as hematopoietic late progenitor *(Ivanova et al. 2002)*, High Output *(Rodriguez-Fraticelli et al. 2020)*, and NoMo (Non-Molecularly Overlapping Output) *(N. K. Wilson et al. 2015)*, in *Usp7***^+/-^**LT-HSCs compared to wild-type controls. In contrast, gene sets linked to HSC quiescence and stemness, including Low output *(Rodriguez-Fraticelli et al. 2020)*, StemScore *(Giladi et al. 2018)* and MolO (Molecularly Overlapping Output) *(N. K. Wilson et al. 2015)* were preferentially enriched in wild-type cells (Figure 4F, Table S3, S8), indicative of less quiescent and more proliferation primed *Usp7***^+/-^** LT-HSCs. Consistent with these findings, Gene Ontology (GO) enrichment analysis of significant DEGs (Table S7) revealed a marked enrichment of cell cycle-related biological processes in *Usp7***^+/-^** LT-HSCs, including chromosome segregation and mitotic progression (Figure S4C). In contrast, GO terms associated with hematopoietic homeostasis and lymphoid differentiation potential were reduced, consistent with the decreased frequencies of MPP4 and CLP populations observed in *Usp7*^+/-^ mice compared with wild-type controls.

Taken together, these results demonstrate that USP7 maintains LT-HSC quiescence and long-term self-renewal, by restraining proliferation-associated programs.

### USP7 maintains LT-HSC identity through control of proteome remodeling

USP7, as a deubiquitinase regulates the stability and the activity of multiple substrates. We therefore performed ultra-low-input (ULI) proteomic analysis to examine the effects of *Usp7* haploinsufficiency on the LT-HSCs’ proteome. After quality filtering, around 6,000 proteins were reliably detected with robust coverage across biological replicates in both genotypes (Figure S4D–G). PCA revealed genotype-dependent clustering with tight grouping of biological replicates, supporting the dataset robustness and indicating global proteomic alterations in *Usp7*⁺^/^⁻ LT-HSCs (Figure S4D). As a control, we assessed the expression of all USP family members in both genotypes. Consistent with previous data *(Cabezas-Wallscheid et al. 2014)* (Figure S1A), we observed expression of the same USP family proteins in LT-HSCs. Notably, USP7 was the only USP whose expression was reduced (∼30%) in *Usp7*^+/-^ cells compared with wild-type controls, indicating the absence of compensatory upregulation by other USP family members (Figure S4G-H). Volcano plot analysis of the proteomic data identified a subset of proteins significantly altered in *Usp7*⁺^/^⁻ LT-HSCs (Figure 4G; Table S9-10). DAVID functional enrichment analysis identified a significant enrichment of GO terms related to metabolic processes, post-translational modifications, and cell-cycle regulation in *Usp7*⁺^/^⁻ LT-HSCs compared with wild-type controls (Figure 4H; Table S11). Consistent with these findings, iBAQ proteomic analysis revealed increased abundance of representative metabolic and cell-cycle regulators in *Usp7*⁺^/^⁻ LT-HSCs (Figure S4G).

Together, these results establish USP7 as a key regulator of LT-HSC homeostasis that restrains proliferative and metabolic pathways.

### USP7 regulates HMGA2 stability

To identify potential post-transcriptionally regulated targets of USP7, we integrated transcriptome and proteome datasets (Figure 5A). Although a significant correlation between mRNA and protein levels was observed, the relationship remained limited (R²=0.27 in both the *Usp7^+/+^* and *Usp7*^+/-^ datasets), underscoring the limited predictive power of transcript abundance for protein levels and highlighting the substantial contribution of post-transcriptional regulatory mechanisms (Figure S5A). A Venn diagram shows the overlap between proteins significantly decreased in *Usp7*^+/-^ LT-HSCs only at the protein level and proteins enriched in LT-HSCs compared with MPP1, as reported by Cabezas-Wallscheid et al., *(Cabezas-Wallscheid et al. 2014)*(Figure 5B). Among the overlapping proteins, we focused on high-mobility group AT hook 2 (HMGA2), as a chromatin-associated architectural transcription factor implicated in HSC self-renewal and proliferation *(Kubota et al. 2024)*. Notably, proteomic analysis revealed a significant reduction in HMGA2 abundance, as well as that of one of its transcriptional targets, in *Usp7*^+/−^ conditions compared with wild-type controls (Figure S4G). We further confirmed the interaction between USP7 and HMGA2 by co-immunoprecipitations followed by western blot in whole-cell lysates from Phoenix cells co-transfected with Flag-USP7 and HMGA2 (Figure 5C; S5B). Immunofluorescence revealed that USP7 and HMGA2 colocalize, supporting their physical interaction observed in co-immunoprecipitation assays (Figure 5D; S5C). Given its deubiquitinating activity, we next asked whether USP7 regulates HMGA2 ubiquitination and protein stability. Consistently, HMGA2 ubiquitination was reduced in USP7-overexpressing cells (Figure 5E), whereas USP7 pharmacological inhibition with XL177A decreased HMGA2 abundance (Figure 5F).

**Figure 5.**
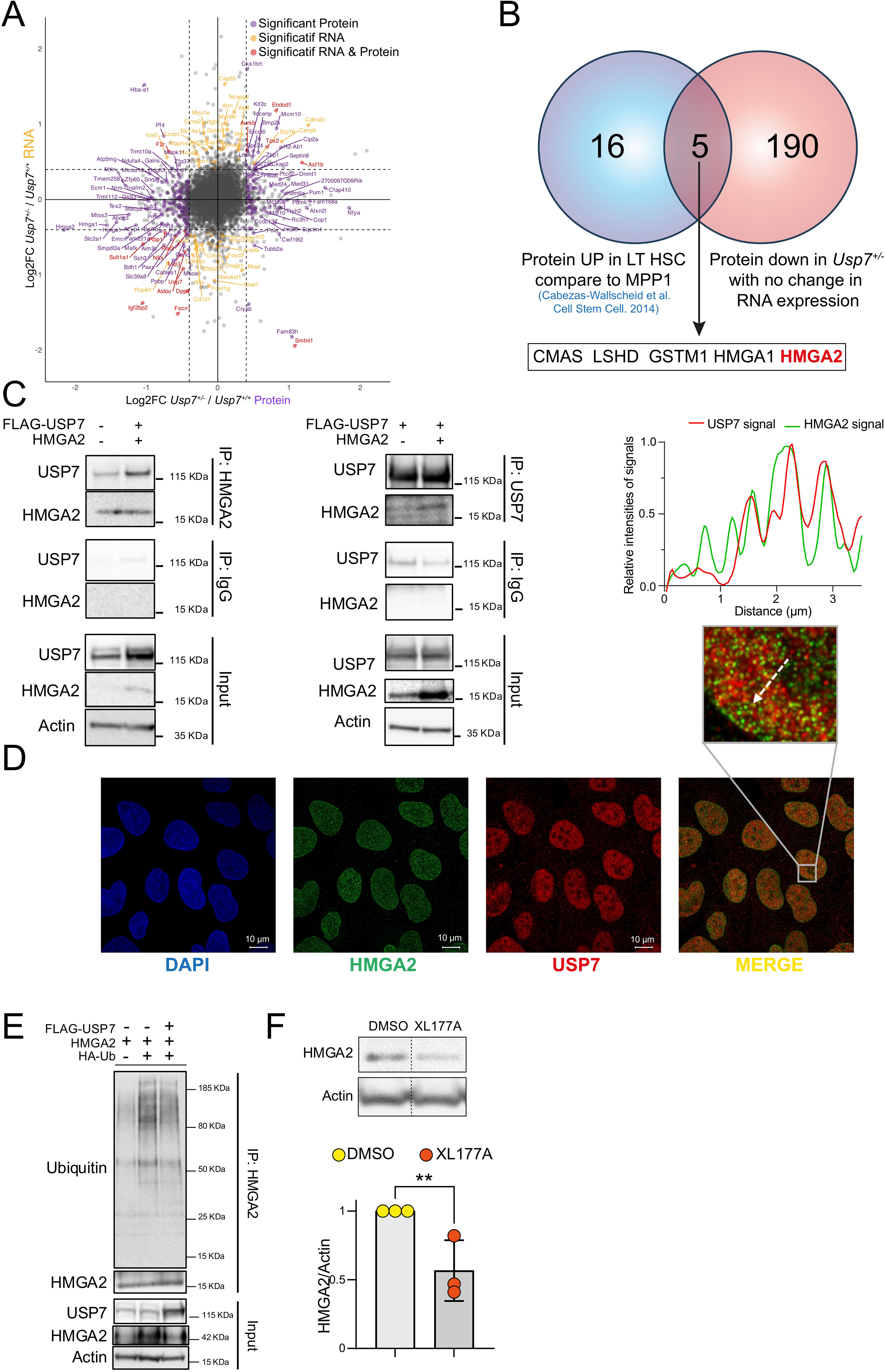
USP7 stabilizes HMGA2 to regulate HSC function. (**A**) Correlation between protein and RNA expression changes. Significant RNA changes (|log₂FC > 0.4|; P-val-adj < 0.05) are shown in orange, and significant protein changes (|log₂FC > 0.4|; q < 0.05) are shown in purple, and concordant RNA-protein alterations are shown in red. (**B**) Venn diagram showing downregulated proteins in *Usp7*^+/-^ compared to *Usp7*^+/+^ LT-HSCs (log 2FC < −0.4; q < 0.05) without downregulation at RNA level in LT-HSCs compared with proteins upregulated in HSCs relative to MPP1 *(Cabezas-Wallscheid et al. 2014)*. Common relevant proteins are indicated and colored. (**C**) Co-immunoprecipitation of USP7 and HMGA2 from transfected Phoenix whole-cell extracts, followed by immunoblotting with antibodies against the indicated proteins. Reciprocal co-immunoprecipitation, confirming the USP7-HMGA2 interaction. (**D**) Immunofluorescence in U2OS cells co-stained with anti-USP7 (red), anti-HMGA2 (green) and DAPI (blue). Images were acquired by confocal microscopy and analyzed using Zen Blue software. (**E)** Ubiquitination of HMGA2. Phoenix cells were co-transfected with USP7-Flag, HMGA2 and HA-Ub constructs for 48h. Sixteen hours prior to harvesting, cells were treated with 20μM MG132. Cell lysates were subjected to immunoprecipitation with control irrelevant immunoglobulin (IgG) or anti-HMGA2 antibody, followed by immunoblotting with anti-Ubiquitin antibody. (**F**) HMGA2 expression 24h following pharmacological inhibition of USP7 with 3μM XL177A in HCT-116 cells. Data represent at least three independent experiments. Results are shown as mean ± SD; ns not significant, **p < 0.01.

Together, these findings demonstrate that USP7 modulates HMGA2 ubiquitination and stability, thereby revealing a post-transcriptional mechanism regulating stem cell quiescence.

## DISCUSSION

Hematopoiesis is a tightly regulated and highly dynamic process that relies on the precise control of long-term hematopoietic stem cell (LT-HSC) fate decisions. Here, we identify USP7 as a critical regulator of LT-HSC dormancy. Within the LSK fraction, MPP6, MPP5, MPP3, and MPP4 populations were markedly depleted in *Usp7*^+/-^ mice, whereas LT-HSCs remained numerically intact but functionally impaired, failing to support long-term reconstitution. These findings highlight that LT-HSC dormancy is not a passive state but a transcriptionally and functionally active program.

Under steady-state conditions, the reduced progenitor (LSK) pool in *Usp7*^+/-^ mice is sufficient to sustain baseline hematopoiesis. However, the functional impairment of LT-HSCs may limit hematopoietic resilience under stress. Supporting this notion, Shan et al. reported that Scl-Cre;*Usp7*^fl/fl^ mice succumbed dramatically earlier than wild-type controls following 5-fluorouracil (5-FU) treatment *(Shan et al. 2026)*, emphasizing the critical requirement for USP7 during regenerative stress. Together, these studies support a model in which USP7 is largely dispensable for steady-state hematopoiesis but essential for maintaining LT-HSC functional integrity under conditions of hematopoietic challenge.

Considering the numerical and functional defects observed in the *Usp7*^+/-^ LSK compartment, we performed single-cell RNA sequencing on sorted LSK cells to define the underlying molecular programs. Analysis of inferred cell-cycle positioning indicated that *Usp7* haploinsufficiency promotes exit from quiescence and increased cell-cycle engagement within the LSK compartment. Consistently, Ki-67 staining revealed a reduced proportion of quiescent (G0) cells among the most immature LSK subsets, including LT-HSCs, MPP6, and MPP5. Given that quiescence is a defining feature of LT-HSCs *(Foudi et al. 2009)*, we next focused our single-cell analyses on this compartment. *Usp7*^+/-^ LT-HSCs exhibited a pronounced shift from dormancy toward an active transcriptional program, characterized by enrichment of the active HSC (aHSC) signature and downregulation of the dormant HSC (dHSC) signature *(Cabezas-Wallscheid et al. 2017)* (Figure 3E). These data support a critical role for USP7 in maintaining transcriptional dormancy in LT-HSCs. Differential expression analysis further revealed a marked reduction of *Meg3*, a lncRNA associated with dormant HSCs *(Cabezas-Wallscheid et al. 2017; Sommerkamp et al. 2019)*, alongside upregulation of the proliferation-associated gene *Hist1h2ap* and *Nupr1*, consistent with increased cell-cycle engagement and altered functional potential. Notably, *Nupr1* deficiency has been reported to enhance HSC engraftment and output *(T. Wang et al. 2022)*, suggesting that its elevated expression in *Usp7*^+/-^ cells may contribute to impaired functional capacity. Collectively, these findings indicate that *Usp7* deficiency disrupts LT-HSC dormancy programs and uncouples quiescence from functional competence, underscoring a critical role for USP7 in maintaining both the quiescent state and functional integrity of LT-HSCs.

To further characterize this phenotype, we performed transcriptomic and proteomic profiling of LT-HSCs. GSEA analyses revealed a significant enrichment of the high-output HSC signature, accompanied by a concomitant downregulation of the low-output HSC signature in *Usp7*^+/-^ LT-HSCs. High-output HSCs are defined by an increased capacity to generate hematopoietic progeny and are associated with enhanced cell-cycle priming and proliferative programs, whereas low-output HSCs exhibit reduced output and are enriched for transcriptional programs linked to quiescence *(Rodriguez-Fraticelli et al. 2020)*. Consistently, signatures associated with dormant HSC states, including the “stemness score” *(Giladi et al. 2018)* and the MolO program *(N. K. Wilson et al. 2015)* were downregulated in *Usp7*^+/-^ LT-HSCs, whereas the NoMo signature, characteristic of active HSCs (aHSC), was enriched *(N. K. Wilson et al. 2015)*. This transcriptional reprogramming was accompanied by enrichment of multiple proliferation-related signatures, including “MYC TARGETS”, previously reported to be associated with aHSCs *(Cabezas-Wallscheid et al. 2017)*. In line with this, proteomic analyses revealed enrichment of proliferative and metabolic programs in *Usp7^+/-^* LT-HSCs. Altogether, these data indicate that *Usp7*^+/-^ LT-HSCs adopt a more activated state, characterized by enhanced proliferative and metabolic activity at the expense of quiescence and differentiation programs, in agreement with previous studies linking HSC activation to metabolic engagement and cell-cycle entry *(Cabezas-Wallscheid et al. 2017; Rettkowski et Cabezas-Wallscheid 2025)*.

To gain deeper insight into the mechanisms governing LT-HSC dormancy, we crossed *Usp7*^+/-^ mice with the H2B-GFP reporter system *(A. Wilson et al. 2008; Foudi et al. 2009)*. In wild-type mice, LT-HSCs retained the highest GFP signal, closely followed by MPP6. Consistently, Sommerkamp *et al*., previously described MPP6 compartment to have a long-term multilineage reconstitution similar to HSCs *(Sommerkamp et al. 2021)*. We also observed intermediate and committed progenitors losing GFP progressively, reflecting established differentiation hierarchies. GFP retention correlated with EPCR expression, confirming that quiescent LT-HSCs preferentially reside in the GFP^+^ EPCR^+^ fraction *(Rabe et al. 2020; Lin et Trumpp 2023)*. In *Usp7*^+/-^mice, GFP retention within LT-HSCs was markedly reduced, with most cells losing detectable signal by six months, whereas a substantial fraction of wild-type LT-HSCs remained GFP^+^. These findings demonstrate that *Usp7* haploinsufficiency depletes the dormant LT-HSC pool, directly compromising stem cell quiescence and providing a mechanistic basis for the functional defects observed under hematopoietic stress.

To better understand how, given that USP7 primarily regulates protein stability through its deubiquitinase activity, we reasoned that functionally relevant targets would be preferentially altered at the protein rather than transcript level. By intersecting proteins selectively deregulated in *Usp7*^+/-^ LT-HSCs at the proteomic, but not at the RNA level with a previously published HSC-enriched proteomic dataset *(Cabezas-Wallscheid et al. 2014)*, HMGA2 emerged as a strong candidate. We demonstrate that USP7 regulates HMGA2 ubiquitination and stability, positioning HMGA2 as a direct downstream effector. Functionally, HMGA2 has been implicated in HSC self-renewal through the Lin28b-let-7 axis during development *(Copley et al. 2013)* and, more recently, has been shown to promote HSC survival and expansion under stress conditions *(Kubota et al. 2024)*, consistent with our observations. Mechanistically, HMGA2 has been shown to directly activate *Igf2bp2* transcription in HSCs, thereby promoting self-renewal programs *(Sun et al. 2022; Cox et al. 2025)*. IGF2BP2, an m6A reader protein, stabilizes key transcripts involved in stem cell maintenance by binding methylated mRNAs and enhancing their stability *(Yin et al. 2022)*. Loss of IGF2BP2 results in impaired HSC quiescence, reduced repopulation capacity, and destabilization of self-renewal-associated transcripts, highlighting its essential role in maintaining HSC function *(Suo et al. 2022; Yin et al. 2022)*. Notably, the coordinated downregulation of *Igf2bp2* at both transcript (Figure 4E) and protein levels (Figure S4G) in *Usp7*-deficient LT-HSCs, supports the existence of a USP7-HMGA2-IGF2BP2 regulatory axis. Given the established role of IGF2BP2 in maintaining RNA stability networks essential for stem cell identity, disruption of this axis is likely to impair LT-HSC quiescence and long-term regenerative capacity in our condition. Together, these findings support a model in which USP7-mediated stabilization of HMGA2 sustains a downstream transcriptional and post-transcriptional program *via* IGF2BP2 to sustain LT-HSC fitness and stress adaptability.

Finally, although PU.1 has recently been implicated as a downstream mediator of USP7 in the regulation of HSC homeostasis, the partial rescue observed upon PU.1 restoration argues against a simple linear pathway *(Shan et al. 2026)*. Rather, these findings, together with our proteomic analyses, reinforce a model in which USP7 functions as a post-translational hub coordinating multiple substrates. These include canonical regulators of cell cycle and mitosis such as MCM10 and INCENP (Figure S4G), as well as potential interactors involved in metabolic regulation such as SLC7A6, PDIA6 (Figure S4G), CMAS, LDHD and GSTM1 (Figure 5B). Through this broad substrate network, USP7 fine-tunes LT-HSC quiescence and self-renewal.

Interestingly, recent studies indicate that *Usp7* deficiency does not impair HSC homing to the bone marrow (*Shan et al. 2026*). However, we show that *Usp7*-deficient LT-HSCs fail to sustain long-term reconstitution and cannot engraft in secondary recipients (Figure 2 and 4C), revealing a profound intrinsic defect in self-renewal. This functional impairment is accompanied by a marked reduction in TGF-β receptor signaling, a key niche-derived regulator of dormancy, as revealed by our RNA-seq analysis of *Usp7*^+/-^ LT-HSC (Figure S4C) *(Yamazaki et al. 2009)*. These findings suggest that USP7 preserves LT-HSC functional integrity not by controlling niche access, but by maintaining responsiveness to dormancy signals, highlighting how intrinsic deubiquitinase activity and extrinsic niche cues converge to regulate stem cell fate, quiescence and stress resilience.

Understanding the mechanisms that enforce HSC dormancy provides key insights into how cancer stem cells survive, particularly in AML. Leukemia stem cells (LSCs) share stem-like features with dHSCs, including self-renewal, differentiation potential, and chemotherapy resistance due to their dormant state, a major driver of AML relapse *(Stelmach et Trumpp 2023)*. In previous work, we showed that USP7 activity contributes to therapy resistance in AML *(Cartel et al. 2021)*. Building on this, our current study identifies USP7 as a critical regulator of LT-HSC dormancy, through HMGA2 regulation. Expression level of HMGA2 have been associated with stem cell-like characteristics and self-renewal capacity in both normal and malignant hematopoietic contexts *(Moison et al. 2022)*. Whether USP7 similarly regulates dormancy in LSCs, as we show here for LT-HSCs, remains an important question for future investigation. Collectively, these findings position USP7 as a shared nodal factor controlling dormancy, stem cell integrity, and therapy response, highlighting its potential as a therapeutic target to mobilize quiescent LSCs and overcome AML resistance and relapse.

## SUPPLEMENTARY FIGURE LEGENDS

**Figure S1.** (**A**) USP family protein expressions were extracted from Cabezas et al. *(Cabezas-Wallscheid et al. 2014)* in HSC and MPP1 (unit: IBAQ, Intensity-Based Absolute Quantification). (**B-C**) *Usp7* gene expression was retrieved from the microarray dataset available in the BloodSpot 3.0 database *(Gíslason et al. 2024)*. (**B**) Mouse hematopoietic populations are shown (extracted from: “mouse normal hematopoiesis system”) (**C**) human hematopoietic populations were analyzed using the same platform from the “normal Human hematopoiesis (DMAP)” dataset. For both species, expression values correspond to log2-normalized Affymetrix microarray intensities. Bars represent mean *Usp7* (mouse) *or USP7* (human) expression, individual points indicate independent samples, and error bars show ± SD. (**D**) Absolute number of total bone marrow cells in 3–5-month-old *Usp7*^+/+^ (n=12) and *Usp7*^+/-^ (n=14) mice. (**E**) Relative expression of USP7 in the total bone marrow of *Usp7*^+/+^(green, n=12) and *Usp7*^+/-^ (blue, n=14) mice analyzed by western blot, 2 independent experiments and (**F**) RT-qPCR (n=3). (**G**) Concatenated representative spectral flow cytometry plots detailing the gating strategy used to define HSPCs and differentiated hematopoietic compartments in the bone marrow of 3-5-month-old *Usp7*^+/+^ (Table S1). (**H-I**) Flow cytometry quantification of mature hematopoietic cell populations (**H**) and of B cell subpopulations (**I**). For all panels, *Usp7*^+/+^ (n=12) and *Usp7*^+/-^(n=14) mice from two independent experiments, except for macrophage and NK cell analyses, which were performed in n = 6 *Usp7*^+/+^ and n = 7 *Usp7*^+/-^ mice from one independent experiment. (**J**) Blood parameters in 3-month-old *Usp7*^+/+^ (n=10) and *Usp7*^+/-^ (n=10) mice. Two independent experiments. Results are shown as mean ± SD; ns not significant, *p < 0.05, **p < 0.01, ***p < 0.001, ****p < 0.0001.

**Figure S2.** (**A**) Workflow of the bone marrow competitive engraftment assay. Whole BM cells from 3-4-month-old CD45.1 wild-type mice and CD45.2 *Usp7*^+/+^or *Usp7*^+/-^ mice were mixed and transplanted into 2-month-old CD45.1 recipient mice. For each genotype, BM cells from each CD45.2 individual donor mice were transplanted into two independent CD45.1 lethally irradiated recipient mice. In total, four CD45.2 donor mice per genotype (*Usp7*^+/+^ or *Usp7*^+/-^) were used. The competitive transplantation was performed using a mixture of CD45.1 (competitor) and CD45.2 (*Usp7*^+/+^ or *Usp7*^+/-^) donor cells (n=8). For secondary transplantation, total BM cells from primary recipients were transplanted into 3-4-month-old CD45.1 lethally irradiated recipient mice (n=8). (**B**) Representative flow cytometry density plots of concatenated bone marrow samples, showing engrafted donor cells (CD45.2⁺) from *Usp7*^+/+^ or *Usp7*^+/-^ mice, competitor bone marrow cells (CD45.1⁺) from wild-type mice, and recipient-derived cells (CD45.1). Plots shown on the left correspond to samples collected before transplantation, and plots on the right display samples collected 1, 2, and 3 months after primary transplantation, as well as 2.5 and 4 months after secondary transplantation. (**C**) Quantification of donor CD45.2⁺ engraftment over time gated on Ter119**⁻** cells. Each green dot represents an individual *Usp7*^+/+^ mouse, and each blue dot represents a *Usp*7^+/-^ mouse. (**D**) Relative donor contribution to myeloid, B-cell, and T-cell lineages before transplantation, at the time of transplantation, after the first, and after the second transplantation. Results are shown as median ± SD; ns: not significant, *p < 0.05, **p < 0.01, ***p < 0.001, ****p < 0.0001.

**Figure S3.** (**A**) Top left: UMAP embedding showing Seurat clusters from unsupervised clustering. Top right: UMAP of cells grouped by source and colored by inferred cell cycle (https://satijalab.org). Bottom left: UMAP after cell cycle correction with clusters re-identified. Bottom right: UMAP depicting cell cycle distribution on the new clusters following correction. (**B**) UMAP of transcriptional clusters from LSK cells of *Usp7*^+/+^ and *Usp7*^+/-^ mice, color-coded by annotated clusters. Data were down-sampled to 20,000 cells per condition. (**C**) Gene set scores for lineage-specific expression signatures derived from scRNA-seq data *(Rodriguez-Fraticelli et al. 2018)* were computed for each annotated cluster. The dot plot shows, for each cluster, the percentage of cells with detectable signature expression (dot size) and the mean signature score (color scale). (**D**) Proportion of the annotated clusters in the LSK population of *Usp7*^+/+^ and *Usp7*^+/-^ mice. (**E**) UMAP projection showing LT-HSCs, ST-HSCs, MPP2, MPP3, and MPP4 populations, identified by supervised classification with the CaSTLe *(Lieberman et al. 2018)* and guided by previously published scRNA-seq data *(Rodriguez-Fraticelli et al. 2018)*. (**F**) Same analysis shown according to genotype. (**G**) Quantification of LT-and ST-HSCs and MPP2-4 fractions (CaSTLe analysis) within each of the identified clusters. (**H**) Inferred *Top2A* (n=61,524) and *Pcna* (n=61,524) expression dynamics with a periodic loess curve plotted according to Tricycle-derived cell cycle positions. (**I**) Violin plots of expression for *Meg3*, *Malat1*, *Meis1*, *Hist1h2ap*, and *Nupr1* in down-sampled LT-HSC cluster (2,000 cells per genotype). P-values were computed using Wilcoxon tests (wilcox.test). (**J**) Representative flow cytometry plots of concatenated samples showing Ki-67 staining in LSKs, LT-HSCs and MPP1-6 subsets from *Usp7*^+/+^ and *Usp7*^+/-^ mice. (**K**) Gating strategy shows the exclusion of dead (Zombie^+^) cells and lineage-positive marker-expressing cells for the identification of LT-HSCs, MPP1-6 subsets, LSK (Lineage⁻ Sca1⁺ c-Kit⁺), GMPs (granulocyte-monocyte progenitors), MEPs (megakaryocyte-erythroid progenitors), and CMPs (common myeloid progenitors). (**L**) Expression of H2B-GFP in all subsets in the bone marrow of *Usp7*^+/+^ and *Usp7*^+/-^ mice after 1, 3 and 6 months DOX chase. Concatenation of all sample data are shown. For each genotype and timepoint, at least 3 mice were analyzed after 1, 3 and 6 months of DOX chase. (**M**) Representative flow cytometry plots and (**N**) quantification of the proportion of EPCR expression within *Usp7^+/+^*and *Usp7^+/-^* LT-HSC cells in GFP-positive or negative populations after 1, 3 and 6 months of DOX chase. Results are shown as mean ± SD; *p < 0.05, **p < 0.01, ***p < 0.001, ****p < 0.0001.

**Figure S4.** (**A**) Gating strategy for LT-HSC sorting. LT-HSCs were defined as Lin⁻Sca-1⁺c-Kit⁺CD48⁻CD150⁺CD34⁻ live singlets. (**B**) Principal Component Analysis (PCA) RNA seq data was performed to visualize sample-to-sample variation. The PCA plot showed clear separation between *Usp7*^+/+^ (WT; n=4) and *Usp7*^+/-^ (HT; n=4) samples. The first two components, PC1 and PC2, explained 40% and 18% of the variability in the gene expression, respectively. (**C**) GO enrichment analysis of genes significantly upregulated (log₂FC > 0.5; P-val-adj < 0.05) or downregulated (log₂FC < −0.5; P-val-adj < 0.05) between *Usp7*^+/+^ and *Usp7*^+/-^ LT-HSCs identifies enriched biological processes. (**D**) Comparison of IBAQ protein expression from MS/MS-based proteomic analysis of *Usp7*^+/+^ and *Usp7*^+/-^ LT-HSCs. Left panel: comparison among *Usp7*⁺^/^⁺ (WT1-4) replicates with each other; Right panel: comparison among *Usp7*^+/-^ (HT1-4) replicates. (**E**) Venn diagram showing protein coverage overlap between four replicates of *Usp7*^+/+^ (WT1-4) and *Usp7*^+/-^(HT1-4) LT-HSCs. (**F**) PCA of proteomic profiles from *Usp7*^+/+^ (WT1-4) and *Usp7*^+/-^ (HT1-4) LT-HSCs. (**G**) Expression of USP7, HMGA2, its transcriptional target IGF2BP2, and selected regulators of metabolism and proliferation, quantified by MS/MS-based proteomic analysis. (**H**) Expression levels of other USP family members in LT-HSCs from *Usp7*^+/+^ and *Usp7*^+/-^ mice, quantified by MS/MS-based proteomics.

**Figure S5.** (**A**) Scatter plots show the correlation between mean RNA expression and mean protein abundance for genes detected in both datasets. RNA levels were quantified as TPM and protein levels as IBAQ, each averaged across four biological replicates per condition (*Usp7*^+/+^ (green) and *Usp7*^+/-^ (blue)). Values were log-transformed as log10(x + 1). Each point represents one transcript/protein. The solid line indicates the least-squares linear regression fit with 95% confidence interval. Pearson correlation coefficient (r), associated p-value, and R² are reported on the plot. (**B**) Phoenix cells were transfected with the indicated plasmids and subjected to immunoprecipitation using anti-HMGA2 antibody or control IgG, followed by immunoblot analysis for ubiquitin. Data are representative of three independent experiments. (**C**) An independent set of images showing USP7 and HMGA2 localization, together with immunofluorescence controls using secondary antibody alone.

## TABLE LEGENDS

**Table S1**. **Spectral Flow Cytometry Antibody Panel for Bone Marrow phenotyping.** The table lists the antibody panel used for spectral flow cytometry–based phenotyping of mouse bone marrow cells. Target antigens, fluorochrome conjugates, antibody clones, dilutions, and manufacturers are indicated. The panel includes markers for hematopoietic stem and progenitor cell identification and lineage characterization, including stem cell markers (e.g., Sca-1, c-Kit, CD150), progenitor-associated markers (e.g., CD34, CD135, CD48, CD16/32), and lineage or differentiation markers (e.g., CD41, CD61, CD105, BP1/CD249 Ly-51, CD127). Related to Fig. 1 and S1G-J; Fig. 2 and S2; Fig. 3F-I and S3J-M; Fig. 4B-C and S4A.

**Table S2**. **Primer sequences and working concentrations.** Primer sequences used in this study are listed together with their final working concentrations in the indicated assays. Primers were designed to ensure specificity and optimal amplification efficiency under the experimental conditions used. Related to Fig.S1F.

**Table S3. Published gene signatures used for projection analyses.** Gene signatures curated from previously published studies and used for projection analyses in this study are listed. These datasets were integrated to enable comparative mapping of transcriptional programs and to define cellular states within the hematopoietic stem and progenitor cell compartment. Related to Fig. 3B, E, Fig. S3C, S3G and Fig. 4F.

**Table S4. Pseudo-bulk differential gene expression analysis (sc-RNAseq) on LT-HSC cluster from *Usp7^+/+^* or *Usp7^+/-^*mice.** List of differentially expressed genes between the indicated groups. The table reports gene symbols, P values (*P_val*) (Wilcoxon rank-sum), average log2 fold change (*avg_log2FC*), and the proportion of expressing cells in each group (*pct.1* and *pct.2*). Adjusted P values (*P_val_adj*) were calculated using the Bonferroni method. Positive log2 fold change values indicate higher expression in group *Usp7^+/-^*relative to group *Usp7^+/+^*. Related to Fig. 3E.

**Table S5. Read count table from RNA-seq data from *Usp7^+/+^*and Usp7^+/-^mice.** Read count table from RNA-seq data aligned to mouse Ensembl-annotated genes generated with featureCounts2. Each row corresponds to one gene, identified by its Ensembl gene ID (Geneid), together with its genomic annotation: chromosome (Chr), start position (Start), end position (End), strand orientation (Strand), and gene/locus length (Length). The following columns indicate the number of reads assigned to each gene across LT-HSC sorted samples, including 4 *Usp7^+/+^* replicates (LT-HSC WT, n1 to n4) and 4 *Usp7^+/-^* replicates (LT-HSC HT, n1 to n4). Related to Fig. 4D.

**Table S6. Normalized Gene expression matrix in LT-HSCs from *Usp7^+/+^*and *Usp7^+/-^* mice**. The table reports normalized gene expression values from RNA-sequencing in long-term hematopoietic stem cells (LT-HSCs). Each row corresponds to one gene, indicated by its gene symbol (gene_name) and Ensembl gene identifier (Ensembl_identifier). The following columns show normalized read counts for each sample, including 4 *Usp7^+/+^*LT-HSC replicates (LT-HSC WT, n1 to n4) and 4 *Usp7^+/-^* LT-HSC replicates (LT-HSC HT, n1 to n4). Related to Fig. 4E-F.

**Table S7. Differential gene expression analysis of LT-HSCs from *Usp7^+/+^* and *Usp7^+/-^* mice**. Differential expression analysis results from RNA-seq data comparing *Usp7^+/+^*and *Usp7^+/-^* samples. Each row corresponds to one gene, indicated by its gene symbol (gene_name) and Ensembl gene identifier (Row.names). The baseMean column represents the mean normalized expression level across all samples. log2FoldChange indicates the expression change between conditions, with positive values corresponding to higher expression in *Usp7^+/-^*LT-HSC and negative values corresponding to lower expression. lfcSE indicates the standard error of the log2 fold change, while stat corresponds to the statistical test value. The pvalue column gives the nominal p-value, and padj corresponds to the adjusted p-value after correction. Related to Fig. 4D and S4C.

**Table S8. Gene set enrichment analysis reveals opposing stemness and proliferative programs.** Gene set enrichment analysis (GSEA) of RNA-seq and single-cell RNA-seq datasets (LT-HSC cluster) showing normalized enrichment scores (NES), nominal p-values, false discovery rate (FDR) q-values and FWER p-value for selected hematopoietic gene signatures. Positive NES indicates enrichment in *Usp7^+/-^* LT-HSCs, whereas negative NES indicates enrichment in *Usp7^+/+^* LT-HSCs. Related to Fig. 3E and Fig.4F.

**Table S9. Quantitative proteomic profiling of LT-HSCs from *Usp7^+/+^*and *Usp7^+/-^* mice**. Protein-level quantification table from mass spectrometry/proteomics analysis. Each row corresponds to one identified protein or protein group, annotated with its UniProt accession (PG.ProteinGroups, PG.ProteinAccessions), gene name (PG.Genes), protein name (PG.ProteinNames), and FASTA header (PG.FastaHeaders). The NrOfStrippedSequences columns indicate the number of unique stripped peptide sequences detected for each protein in the different LT-HSC samples, including *Usp7^+/+^*(WT) and *Usp7^+/-^* (HT) replicates. The Quantity columns report the protein abundance values quantified in each sample, while the IBAQ columns correspond to intensity-based absolute quantification values. Related to Fig. 4G, S4D-H.

**Table S10. Differential proteomic comparison between *Usp7^+/+^*and *Usp7^+/-^* LT-HSCs.** Differential protein abundance analysis comparing USP7 heterozygous and wild-type LT-HSC samples. Each row corresponds to one protein or protein group identified by its UniProt accession (ProteinGroup, UniProtIds) and associated gene name (Genes). The Comparison (group1/group2) column indicates the tested contrast, here *Usp7^+/-^*(HT) / *Usp7^+/+^* (WT). AVG Log2 Ratio represents the average log2 fold-change in protein abundance, with positive values indicating higher abundance in the *Usp7^+/-^* condition. Pvalue corresponds to the nominal statistical significance, while Qvalue indicates the multiple-testing-adjusted significance. The # of Ratios column reports the number of quantitative peptide or protein ratios used for the comparison. Protein annotations are provided in the ProteinDescriptions and ProteinNamescolumns. Related to Fig. 4G, S4D-H.

**Table S11. DAVID enrichment analysis of proteins upregulated in LT-HSCs *Usp7^+/-^* compared to *Usp7^+/+^*.** Functional enrichment analysis was performed on upregulated proteins in *Usp7^+/-^*identified by quantitative mass spectrometry using David tool from NIH (www.davidbioinformatics.nih.gov) Significantly enriched Gene Ontology (GO) Biological Process terms are shown for the top cluster. For each term, the number of proteins in the input list (Count), total list size, population hits, and total population size are indicated, together with the percentage of proteins associated with each term, enrichment fold, nominal p value, and Benjamini-adjusted p value. Related to Fig. 4H.

## METHODS

### Mice

Heterozygous *Usp7* mice floxed *(Kon et al. 2011)* were kindly provided by Professor Wei Gu (Columbia University, NY, USA) and backcrossed with Zp3-Cre strain (kindly provided by Dr J-C. Guéry) on a C57BL/6J background, their genotypes were verified by PCR (Table S2). C57BL/6J Ly5.2, (CD45.2) mice were purchased from Charles River Laboratories (France) and Ly5.1 (CD45.1) congenic mice were initially obtained from The Jackson Laboratory (Bar Harbor, ME) or bred in-house. CD45.1/2 were generated by crossing H2B-GFP transgenic mice (H2B-GFP^tg^) express the histone H2B-GFP^tg^ fusion protein under the control of a tetracycline-responsive regulatory element *(A. Wilson et al. 2008)* were obtained from The Jackson Laboratory (Bar Harbor, ME). H2B-GFP^tg^ mice were backcrossed with heterozygous *Usp7* mice. Genotypes were confirmed by PCR analysis (Table S2), and all experiments were performed using H2B-GFP^tg^ hemizygous offspring. All animals were housed in pathogen-free conditions (Anexplo US006 CREFRE, Toulouse, France) in accordance with the European Directive 2010/63/EU and the French Institutional Guidelines for animal handling. Mice were handled according to protocols approved by the Regional Ethical Committee (#A31555010). Mice were backcrossed into the C57BL/6J background for more than 10 generations.

### Cell Lines

HEK 293T/17 (ATCC, cat. # CRL-11268), Phoenix-AMPHO (ATCC, cat# CRL-3213), HCT-116 (ATCC, cat#CCL247) and U2OS (ATCC, cat#HTB-96) cells were grown in Dulbecco’s modified Eagle medium Glutamax (Life Technology) supplemented with 10% FBS, 100 IU/mL penicillin, and 100 μg/mL streptomycin (Life Technology).

### Flow Cytometry

#### Phenotypic analysis

*Phenotypic analysis of hematopoietic compartments.* Mouse bone marrow (BM) cells (1×10^7^ per sample) were filtered through a 40-µm cell strainer to remove aggregates, and red blood cells were lysed using ACK Lysing buffer (gibco, #A10492-01) for 3 min. Cells were then incubated with Zombie Aqua viability dye (Biolegend, #423102 1:800) in 100 µL PBS 1×, washed, and stained with the antibody cocktail (Table S1A) for 30 min at 37°C in PBS 1× supplemented with 2% FBS (Gibco, #A5256701) and 2 mM EDTA (Sigma-Aldrich #E5134-250G), containing 5% True-Stain Monocyte Blocker™ (BioLegend, #426102) and 10% Brilliant Stain Buffer Plus (BD Horizon™, #566385). Cells were washed in PBS 1×/2% FBS/2 mM EDTA prior to acquisition. Data were acquired on a Cytek Aurora 5L spectral flow cytometer (Cytek Biosciences, Fremont, CA, USA). Raw data were unmixed using SpectroFlo software (v3.3.0; Cytek Biosciences) and then analyzed using OMIQ software (OMIQ Inc., Santa Clara, CA, USA). Antibodies used are listed in Table S1A.

#### UMAP

For the whole hematopoiesis representation, 1.2.10^6^ cells from 6 *Usp7^+/+^* samples were used. Dimensionality reduction was performed using the UMAP algorithm (uwot package) on these 25 markers: CD16/32, CD4, CD11b, CD2, CD61, Ly-6G, CD150 (SLAMF1), F4-80, CD19, CD48, CD8 alpha, Sca-1, CD34, CD335 (NKp46), CD41, CD105, CD201, CD23, CD135 (FLT3), CD117 (c-kit), CD45R (B220), CD127 (IL-7Rα), CD249, IgK-L, and Ly-6C, with the following parameters: n_neighbors = 30, min_dist = 0.5, metric = cosine. Annotated populations, based on the supervised gating, were overlaid.

For the HSPC-focused analysis, cells were pre-gated on hematopoietic stem and progenitor cells (Lineage⁻ Kit^+/low^). UMAP was computed on a restricted panel of 10 functionally relevant markers for HSPC identity: CD16/32, CD150 (SLAMF1), CD48, Sca-1, CD34, CD105, CD201, CD135 (FLT3), CD117 (c-kit), and CD127 (IL-7Rα), using identical parameters to the whole hematopoiesis Annotated populations (LT-HSC, MPP1-6, CMP, GMP, MEP, CLP) based on the supervised gating, were overlaid. Individual marker expression plots were generated for each of the 10 markers using a common min-max color scale gradient.

#### Cell sorting

For cell sorting, mouse bone marrow cells from *Usp*7^+/+^or *Usp*7^+/-^ mice were harvested by flushing, and lineage-positive cells were depleted using the Lineage Cell Depletion Kit (STEMCELL Technologies, #19856A) according to the manufacturer’s instructions. The enriched lineage-negative fraction was stained with antibodies for flow cytometry listed in Table S1B. The Lineage-negative (Lin-), Sca-1+ c-Kit+ (LSK) or LT-HSC population was sorted using BD FACSAria™ II cell sorter (BD Biosciences, San Jose, CA, USA) and then used for reconstitution assays or single-cell RNA sequencing (LSK) and reconstitution assays, RNAseq or proteomics (LT-HSC).

### Bone marrow transplantation and competitive reconstitution assays

For the generation of 50/50 bone marrow chimeras, a total of 8.10⁶ bone marrow (BM) cells were transplanted per recipient. Briefly, 4.10⁶ total BM cells from *Usp7^+/+^* and *Usp7^+/-^* CD45.2 (3-month-old) donor mice were mixed with 4.10⁶ total BM cells from CD45.1 congenic donors, and the cell suspension was injected intravenously into lethally irradiated (9 Gy) CD45.1 recipient mice (2-month-old). For secondary transplantations, 4.10⁶ total BM cells were harvested from primary recipients and injected into lethally irradiated secondary CD45.1 mice (3-month-old).

For LSK transplantation assays, 5,000 sorted LSK (Lineage⁻ Sca-1⁺ c-Kit⁺) cells from 2-month-old wild-type or heterozygous *Usp7* CD45.2 mice were mixed with 5,000 LSKs from wild-type CD45.1 donors (2-month-old) and 5.10⁵ total BM support cells (3-month-old CD45.1/2 mice). The mixture was transplanted into lethally irradiated (9 Gy) CD45.1/2 recipient mice aged 3-4 months. For secondary transplantations, 4.10⁶ total BM cells were harvested from primary recipients and injected into lethally irradiated secondary CD45.1/2 mice (3-month-old).

For LT-HSC transplantation assays, 250 sorted LT-HSCs from *Usp7*^+/+^ and *Usp*7^+/-^ CD45.2 mice (3-5-month-old) were mixed with 1.10⁶ BM support cells from CD45.1/2 mice (2.5-month-old) and transplanted into lethally irradiated (2 × 4.5 Gy) CD45.1 recipients (2-month-old).

Reconstitution and donor chimerism were monitored by flow cytometry using the markers listed in Table S1C. Data acquisition was performed on a BD Fortessa X-20 cytometer (BD Biosciences, San Jose, CA, USA), and analyses were performed using OMIQ software. To assess the reconstitution of hematopoietic lineages in the total bone marrow competition assay, B lymphocytes were identified based on CD19 positivity, myeloid cells were defined as CD11b-positive cells, and T lymphocytes were identified as CD3e-positive cells.

### Cell cycle analysis

Bone marrow cells were stained for surface markers (Table S1D), fixed and permeabilized using Transcription factor Buffer Set (BD Pharmingen, #562574), according to the manufacturer’s instructions. Intracellular staining for Ki-67 was then performed using the Perm/Wash solution (BD Pharmingen). Samples were acquired on a BD Fortessa X-20 cytometer (BD Biosciences, San Jose, CA, USA), and data were analyzed using OMIQ software.

### *In vivo* cell-division assay

To monitor proliferation dynamics and quiescence of hematopoietic stem cells (HSCs), *Usp7*^+/+^ and *Usp7*^+/-^ mice were crossed with homozygous H2B–GFP^tg^ mice to generate heterozygous H2B–GFP^tg^ and *Usp7*^+/-^;H2B–GFP^tg^ offspring, respectively. To induce expression of the H2B-GFP transgene, 8-16-week-old mice received a single intraperitoneal injection of doxycycline (40 mg/kg; Sigma-Aldrich, #D9891) to initiate H2B-GFP labeling, followed by doxycycline administration in the drinking water (2 g/L) supplemented with sucrose (Sigma-Aldrich, S9378) for 6 weeks as previously described *(A. Wilson et al. 2008; Foudi et al. 2009)*. Following doxycycline withdrawal, mice were analyzed at 0, 1, 3 and 6 months to track GFP fluorescence in all compartments, identified by phenotypic staining (Table S1E) and acquired on a Cytek Aurora 5L spectral flow cytometer (Cytek Biosciences, Fremont, CA, USA). GFP quantification was performed using SpectroFlo and a custom R pipeline. At each time point, *Usp7*^+/+^ and *Usp7*^+/-^ cells were ranked from lowest to highest GFP fluorescence intensity, and the area under the curve (AUC) was subsequently calculated. Green and blue dotted lines indicate the standard error of the mean (SEM) for *Usp7*^+/+^ and *Usp7*^+/-^ AUC, respectively. The proportion of GFP-retaining cells is indicated for each time point.

### qPCR analysis

For qPCR, total RNA of 500 000 cells were isolated using RNeasy Plus Micro Kit (Qiagen #74034) and reverse-transcribed using SuperScript VILO cDNA Synthesis (ThermoFisher #11754050) according to the manufacturer’s guidelines. For qPCR analysis, KAPA SYBR FAST (Sigma KK4604) was used on a Real-Time PCR System (Stepone). *Usp7* RNA expression was normalized to *Gusb* housekeeping gene expression and presented as relative quantification (ratio = 2exp-DDCT). Primer sequences are available in the supplementary Information File (Table S2)

### scRNA-seq

#### Purification of LSK

LSK were sorted as previously described and using BD FACSAria™ II cell sorter and gently resuspended at 1,000 cells/μL in PBS 1×/2% FBS/2 mM EDTA.

#### Single-Cell Library Preparation and Sequencing

Between 10,000 and 18,000 cells per sample were encapsulated using the 10X Chromium controller (10X Genomics) following the Chromium Next GEM Single-Cell 3ʹ User Guide v3.1 (10X Genomics). Briefly, cells were encapsulated into oil droplets with barcoded Gel Beads and reagents to convert mRNA into complementary DNA (cDNA). cDNAs were then amplified, fragmented, and Illumina adapters were added during ligation. After performing a last dual index PCR, library quality was assessed using DNF-474 HS NGS kit for Fragment Analyzer (Agilent) and frozen before quantitation and sequencing. Barcoding was done on three levels: cell barcodes allow attribution of each sequence read to its cell of origin; unique molecular identifiers upstream to poly(d)T primers allow tagging of each original molecule to avoid amplification bias; and index allows pooling of different samples. scRNA-seq libraries were quantified by a fluorimetric method using Qubit HS DNA kit (Qubit 4.0, Thermofisher Scientific) and by qPCR using the KAPA library quantification (Roche Diagnostics). The libraries were pooled at equimolar concentrations and sequenced with 1% phiX using an Illumina NovaSeq6000 sequencer, at a sequencing depth of 9,000 to 23,000 reads per cell. Sequencing was configured paired-end at 28 cycles for the first read, 91 cycles for the second one, and 8 cycles for each index.

#### Data Analysis

The preprocessing of raw sequencing data was performed using the 10x Genomics Cell Ranger pipeline v7.1.0 *(G. X. Y. Zheng et al. 2017)*. Reads have been aligned to the mouse reference genome GRCm38 M23 with cellranger count. The data were then analyzed in R v4.2.3, primarily using the Seurat v5.0 package *(Hao et al. 2021)*. Cells with fewer than 200 unique genes (too little information), more than 5,000 unique genes (duplicates), and more than 5% of mitochondrial DNA (dead cells) are sorted out and the four samples are merged.

For the visualization part, gene counts were normalized with a regularized negative binomial regression, using the SCT normalization workflow from the Seurat package *(Hafemeister et Satija 2019)* with a regression on the proportion of mitochondrial reads and on the cell cycle state (inferred with Seurat’s CellCycleScoring function). This normalization allows us to visualize the cells with or without the effect of the cell cycle using uniform manifold approximation and projection (UMAP) dimensionality reduction, using 15 dimensions and spread = 3.

Cell clusters were then identified using Louvain clustering within the Seurat workflow, with the resolution parameter set to 1.2. The annotation of clusters was performed using the Seurat AddModuleScore function with signatures from literature *(Rodriguez-Fraticelli et al. 2018; Table S3)*. Cluster 21 which expressed both the pDC and pMo signatures, was reclassified with a resolution of 0.03 for the final annotation.

For the inference of cell cycle, we applied the Tricycle package version 1.18.0 *(S. C. Zheng et al. 2022)*. To determine the cell cycle stage, we grouped cells between 1.75 π and 0.1 π as G1/G0, those between 0.1 and π as S, and those between π and 1.75 π as G2/M.

Cells annotated as LT-HSC were subset, the RNA assay was normalized (NormalizeData(), Seurat v5.0), RNA layers were joined (JoinLayers()), and differential expression comparing *Usp7^+/+^* versus *Usp7^+/-^* was computed with the FindMarkers() function in Seurat (Table S4) and the resulting gene-level statistics were used to rank genes for enrichment testing in Gene Set Enrichment Analysis (GSEA; v4.3.3; *Subramanian et al. 2005*) on the active HSC (aHSC) and dormant HSC (dHSC) gene signatures *(Cabezas-Wallscheid et al. 2017)*.

We used CaSTLe (Classification of Single Cells by Transfer Learning), a supervised method that labels cells in a scRNA-seq dataset using knowledge learned from related reference datasets *(Lieberman et al. 2018)*. As a source dataset, we selected a published scRNA-seq dataset generated from FACS-isolated HSPCs *(Rodriguez-Fraticelli et al. 2018)*. Cells in this dataset (approximately 2000 per type) are classified into five subsets: LT-HSC (Lin- Sca1+ Kit+ CD135- CD150+ CD48-), ST-HSC (Lin- Sca1+ Kit+ CD135- CD150- CD48-), MPP2 (Lin- Sca1+ Kit+ CD135- CD150+ CD48+), MPP3 (Lin- Sca1+ Kit+ CD135- CD150- CD48+), and MPP4 (Lin- Sca1+ Kit+ CD135- CD150- CD48+ CD135+).

### Bulk RNA-seq

#### RNA Extraction and Library Preparation

Total RNA was extracted from at least 1,000 LT-HSC pooled from a minimum of 5 mice using the Qiagen RNeasy Micro Kit (Cat. No. 74034) according to the manufacturer’s instructions (n=4, *Usp7*^+/+^ and *Usp7*^+/-^, 2 independent experiments). RNA-seq library preparation was performed using the Illumina TruSeq Stranded mRNA Low Sample protocol, starting from 300 ng of total RNA. Cluster generation and sequencing were conducted on an Illumina NextSeq 500/550 High Output Kit v2.5 with a read length of 2 × 75 nucleotides. Paired-end sequencing was performed on an Illumina NextSeq 500.

##### Read Processing, Alignment, and Gene Quantification

The average number of reads per sample was 5 million. Raw RNA-seq reads were first trimmed for adapters and low-quality bases using fastp. Cleaned reads were then aligned to the mouse reference genome GRCm39 (GENCODE Release M36) using STAR v2.7.10b *(Dobin et al. 2013)*. Gene-level counts were generated with featureCounts2 *(Liao et al. 2014)* using gene annotations from GENCODE Release M36 (GRCm39) provided by the GENCODE project.

#### Preprocessing and analysis

Raw count data were filtered to remove lowly expressed genes. Genes were retained only if they showed at least 5 reads in a minimum of two samples, ensuring that low-expression features did not contaminate downstream analyses. Variance-stabilized data were obtained using the regularized log transformation (rlog) implemented in DESeq2 *(Love et al. 2014)*.

Batch effects were corrected for by including batch as a covariate in the DESeq2 design formula (∼ Batch + condition). In addition, batch effects were corrected for exploratory analyses and visualization using the removeBatchEffect function from the limma package *(Ritchie et al. 2015)*. Principal component analysis (PCA) was performed on rlog-transformed data after batch correction.

Differential expression between *Usp7*^+/+^ and *Usp7*^+/-^ conditions was assessed using the Wald test implemented in DESeq2. P-values were adjusted for multiple testing using the Benjamini-Hochberg false discovery rate (FDR) procedure *(Benjamini et Hochberg 1995)*. Independent filtering was disabled to retain adjusted p-values for all genes. Genes with an adjusted p-value (P-val-adj) < 0.05 and an absolute log2 fold change > 0.5 were considered significantly differentially expressed.

Ensembl gene identifiers were mapped to official gene symbols using the biomaRt package with the Ensembl mouse dataset mmusculus_gene_ensembl *(Durinck et al. 2009)*. Gene symbols were subsequently converted to Entrez Gene identifiers using the org.Mm.eg.db annotation package (Carlson, Bioconductor) for Gene Ontology (GO) analysis.

GO enrichment analysis was performed separately for upregulated and downregulated genes using over-representation analysis using the enrichGO function of the clusterProfiler package *(Wu et al. 2021)*. Enrichment analysis was based on Gene Ontology Biological Process annotations provided by the Gene Ontology Consortium. P-values were adjusted for multiple testing using the Benjamini-Hochberg method and GO terms with a P-val-adj < 0.05 were considered significantly enriched.

GSEA was performed using GSEA v4.3.3. Gene signatures used in this study are listed in Table S3. Analyses employed 1,000 permutations (gene_set permutation type), with all other parameters set to default. Results are shown in Table S8.

### Proteomics

#### Sample Preparation

For low-input proteomic analysis, 1,000 sorted LT-HSC pooled from a minimum of 4 mice (4 x *Usp7*^+/+^ and 4 x *Usp7*^+/-^) were collected. The cell pellets were lysed in 6 μL of lysis buffer (100 mM pH 8.5 Triethylammonium bicarbonate (TEAB, T7408 Sigma-Aldrich), 0.2% n-dodecyl-β-D-maltoside (DDM, D4641 Sigma-Aldrich), 10 mM Tris(2-carboxyethyl) phosphine (TCEP, 646547 Sigma-Aldrich), 40 mM 2-Chloroacetamide (CAA, 22790 Sigma-Aldrich)) through three freeze-thaw cycles. Protein digestion was performed by adding 400 ng of trypsin (V5111, Promega) per sample and incubating overnight at 37°C in a humid chamber. The following day, trypsin activity was halted by adding 1% formic acid, and 2% acetonitrile was used to solubilize the peptides allowing for the direct injection of 6 μL ready for MS analysis.

#### Liquid Chromatography–Tandem Mass Spectrometry

The LC-MS/MS analysis was conducted using a timsTOF Ultra mass spectrometer equipped with a CaptiveSpray source (Bruker) and coupled with a nanoElute® 2 nanoflow liquid chromatography system (Bruker). Samples were loaded at 800.0 bar on a PepSep® ULTRA (25 cm x 75 µm x 1.5 µm) C18 HPLC column (Bruker) and equilibrated in 98% solvent A (H2O, 0.1% FA) and 2% solvent B (ACN, 0.1% FA). Peptides were eluted using a two to 4% gradient of solvent B during 1 min, then a 4 to 20% gradient of solvent B during 19 min, followed by a 20 to 30% gradient of solvent B during 5 min and finally a 30 to 95% gradient of solvent B during 1 min all at 250 nl/minute flow rate.

The instrument method for the timsTOF Ultra was set up in a data-independent acquisition (DIA) scan mode that uses parallel accumulation-serial fragmentation (PASEF®) technology (dia-PASEF®). The mass range for MS scans was set up to 100-1700 m/z, whereas the mobility (1/K0) range was set up to 0.64-1.45 Vs/cm² with a 100.0 ms ramp time. The number of MS/MS windows was set up to 24 with a width of 25 Da spanning from 400.0 to 1000.0 Da.

#### Data Analysis

After HTRMS conversion, acquired raw data were analyzed using Spectronaut® v19 (Biognosys, *(Bruderer et al. 2015)*) using the directDIA™ library-free workflow and the Mus musculus reference proteome database (54739 entries, download date July 24, 2018). All searches were performed with oxidation of methionine and protein N-terminal acetylation as variable modifications and cysteine carbamidomethylation as fixed modification. Trypsin was selected as protease allowing for up to two missed cleavages. The maximum and minimum peptide length were set to 52 and 7 amino acids, respectively. The false discovery rate (FDR) for peptide and protein identification was set to 0.01.

#### Statistical analysis of proteomics data

The statistical analysis of the proteomics data was performed as follows: four biological replicates were acquired per condition. Protein quantitative values were measured using Spectronaut (PG.Quantity). To identify proteins that were more abundant in one condition than in another, the quantified values from both conditions were compared. Only proteins with at least two quantitative values in one of the two conditions compared were retained for further statistical analysis to ensure a minimum of replicability. The remaining values were then log2-transformed and normalized by median centering within each condition. Next, missing values of the remaining proteins were imputed using the impute.mle function of the R package imp4p *(Gianetto et al. 2020)*. Statistical testing was conducted using limma t-tests thanks to the R package limma *(Ritchie et al. 2015)*. The FDR control was performed using an adaptive Benjamini–Hochberg procedure on the resulting p-values thanks to the function adjust.p of R package cp4p using the robust method to estimate the proportion of true null hypotheses among the set of statistical tests *(Giai Gianetto et al. 2016)*. The proteins associated with an absolute Log2FC superior to 0.40 and P-val-adj inferior to an FDR (q-value) level of 0.05 have been considered as significantly differentially abundant proteins. All the proteins identified by MS have been used as background for the enrichment tests.

For functional enrichment analysis, proteins that were significantly upregulated in the *Usp7*^+/-^ condition were selected. Analysis was performed using the DAVID bioinformatics tool, using the following annotation categories: GOTERM_BP_ALL, UP_SEQ_FEATURE, UP_KW_PTM, UP_KW_BIOLOGICAL_PROCESS, KEGG_PATHWAY and UP_KW_DOMAIN and considering only terms with a Benjamini-corrected p-value < 0.1 in each cluster. The top three clusters were presented.

### Western Blot and immunoprecipitation

Cells were lysed in NuPAGE LDS Sample Buffer supplemented with NuPAGE Sample Reducing Agent (Life Technologies) and heated for 5 minutes at 95°C. Then proteins were quantified by Micro BSA Protein assay kit (Thermo Fisher Scientific, cat. # 23235) and separated using 4% to 12% gradient polyacrylamide SDS–PAGE gels (Life Technologies) and electrotransferred to 0.2 μm nitrocellulose membranes (Amersham). After blocking in Tris-buffered saline with 0.1% Tween and 5% BSA, membranes were blotted overnight at 4°C with the appropriate primary antibodies. Primary antibodies were detected using the appropriate horseradish peroxidase–conjugated secondary antibodies. Immunoreactive bands were visualized by enhanced chemiluminescence (Thermo Fisher Scientific, cat. # PI32209) with a Pxi camera (Syngene) using GeneSys software (Syngene). Protein levels were quantified using Fiji or genesis software and normalized to nonvariable proteins such as Actin. Primary antibodies used were Actin (cat. #MAB1501) was purchased from Millipore; USP7 (Bettyl, cat. #A300-033A) was purchased from EuroMedex; HMGA2 (Cell signaling, #5269S), and Ubiquitin (Cell signaling, #3936S) were purchased from Cell Signaling Technology.

For immunoprecipitation, approximately 15.10^6^ Phoenix cells overexpressing HMGA2, USP7, HA-ubiquitin or control vector were lysed in an ice-cold lysis buffer (20 mM Tris-HCl pH 7.4, 100 mM NaCl, 0.5% NP-40, Phosphatase Inhibitor cocktails 2 (Merck, #P5726) and 3 (Merck, #P0044) supplemented with protease inhibitors (Roche #05892970001). Alternatively, for ubiquitination assays, cells were treated with 20 µM MG132 (T2154, tebubio) for 16 h prior to lysis to inhibit proteasomal degradation and allow accumulation of ubiquitinated proteins. Lysates (500 µg −1mg total protein) were incubated overnight at 4 °C with 2 µg of anti-HMGA2 antibody (Cell Signaling Technology, cat. #5269S), or anti-USP7 antibody (Abcam, ab106931) or normal rabbit IgG (Cell Signaling Technology, #2729) as control, followed by incubation with Protein A/G magnetic beads (Thermo Fisher Scientific, cat. #88803) for 1 h at 4°C. Beads were washed three times in the lysis buffer, and bound proteins were eluted by boiling in NuPAGE LDS Sample buffer containing reducing agent for subsequent immunoblot analysis.

### Immunofluorescence

U2OS cells were seeded on Ibidi (ibidi GmbH, #80286) and grown to ∼70% confluency. Cells were fixed with 4% PFA for 10 min, permeabilized and blocked with PBS - BSA 3% - 0.1% Triton X-100 for 30 min. Primary antibodies (α-USP7 Thermofisher #MA5-31515 and α-HMGA2 Abcam #ab251468) were applied for 1hour at room temperature in blocking solution, followed by fluorophore-conjugated secondary antibodies for an additional hour at room temperature. Nuclei were counterstained with DAPI 1µM (#1050-A Euromedex). Images were acquired on a confocal microscope (Zeiss LSM980 AIRYSCAN) under consistent settings and processed using Zen Blue software (Zeiss, version 3.11). Colocalization between signals was obtained using Zen Blue profile tool.

To facilitate the visual assessment of spatial overlap and colocalization between both fluorescence signals, intensity line profiles were normalized using a Min-Max scaling approach. This transformation rescales the raw fluorescence intensity values of each channel to a relative range between 0 and 1.

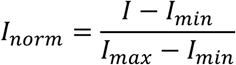

### Expression Vectors

Human USP7 WT ORFs were cloned by Gordon Peters and Goedele Maertens *(Maertens et al. 2010)*; Addgene plasmid #46751. Ubiquitin tagged HA was cloned by Edward Yeh’s laboratory *(Kamitani et al. 1997)*; Addgene # 18712. Mouse HMGA2 ORFs were cloned in pcDNA3.1 (gift from David Bartel, *(Mayr et al. 2007)*); Addgene plasmid # 14789.

### Statistical analysis

Statistical analyses were conducted using GraphPad Prism software v10.6.0. For *in vitro* studies, statistical significance was determined by the two-tailed unpaired Student t test with Welch correction, or paired t test or ratio paired t test when appropriate. For in vivo studies, statistical significance was determined by the nonparametric Mann–Whitney test. Unless otherwise indicated, all data represent the mean ± SD from at least three independent experiments. *, P < 0.05; **, P < 0.01; ***, P < 0.001.

## DATA AVAILABILITY STATEMENT

Further information, resources, and reagents are available upon request. Inquiries should be directed to and will be fulfilled by Antoine Nouhaud and Christine Didier.

## AUTHOR CONTRIBUTIONS

AN, AD, CD designed the study, performed and analyzed the experiments and wrote the manuscript. CD, MP, ED, CB, MM, BG, GA and LL, reviewed the data and the manuscript. CD acquired the funding, CD, AN, AD, MB, QR, HS, NP, SL, DM, PE and SH performed and analyzed experiments, CD conceived and supervised the project and wrote the manuscript. All authors contributed to the final draft.

## ACKNOWLEDGMENTS

We acknowledge Manon Farcé, Fabien Porta, Emeline Sarot, Carine Valles, and Clémentine Dechamps from the Cancer Research Center of Toulouse (INSERM U1037), for their assistance with cell sorting, experimental procedures, and single-cell RNA-seq data preprocessing. We thank all members of mice core facilities (UMS006, ANEXPLO, Inserm, Toulouse) for their support and technical assistance. We thank Dr Frédérique Ruf-Zamojski for critically reading the manuscript and providing helpful comments. This study was supported by institutional grants from the Institut National de la Santé et de la Recherche Médicale (INSERM, France), from the Centre National de la Recherche Scientifique (CNRS, France), from Société Française de lutte contre les Cancers de l’Enfant and the Fondation ARC pour la Recherche sur le Cancer (ARC-PJA2022070005312) and the CARe graduate school. The team is supported and Labeled by the Ligue Nationale Contre le Cancer and the associations : “Laurette Fugain”, “Les 111 des Arts”, “Cassandra” and “Constance la petite guerrière astronaute”. The team is a member of OPALE Carnot Institute.

