## Supplementary figures and images for "USP7 maintains hematopoietic stem cell dormancy and function by stabilizing HMGA2"

### Supplemental Fig S1

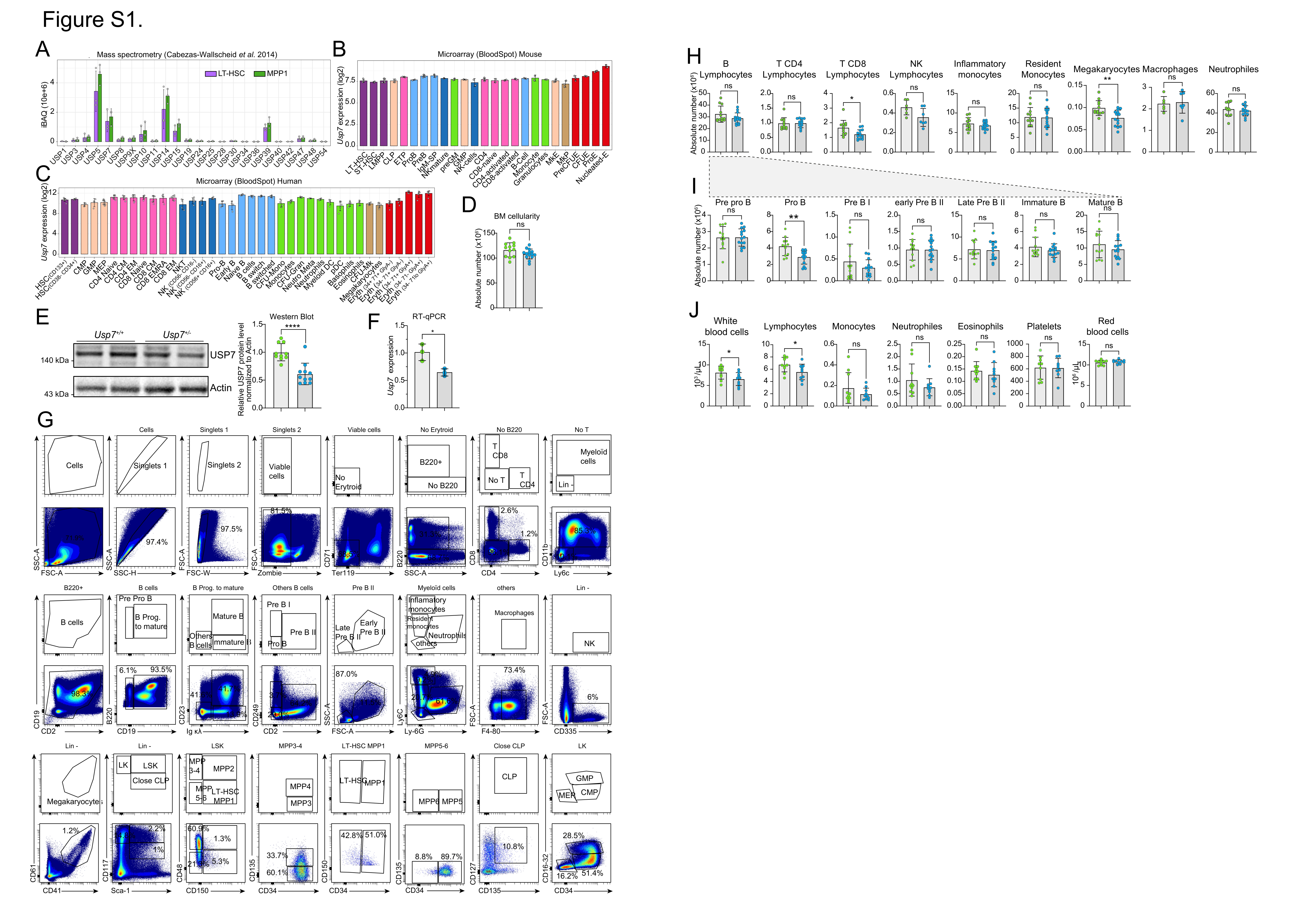

### Supplemental Fig S2

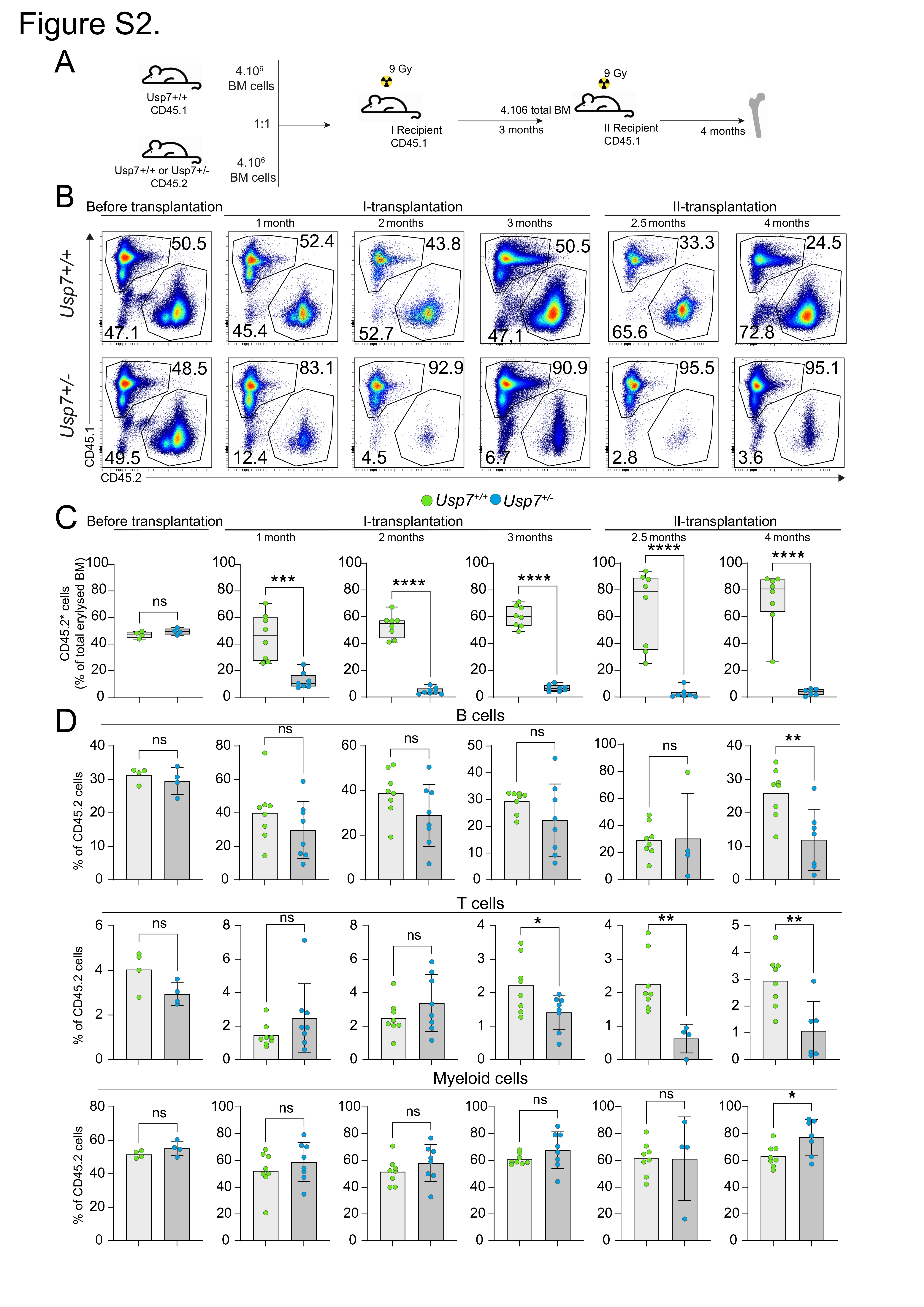

### Supplemental Fig S3

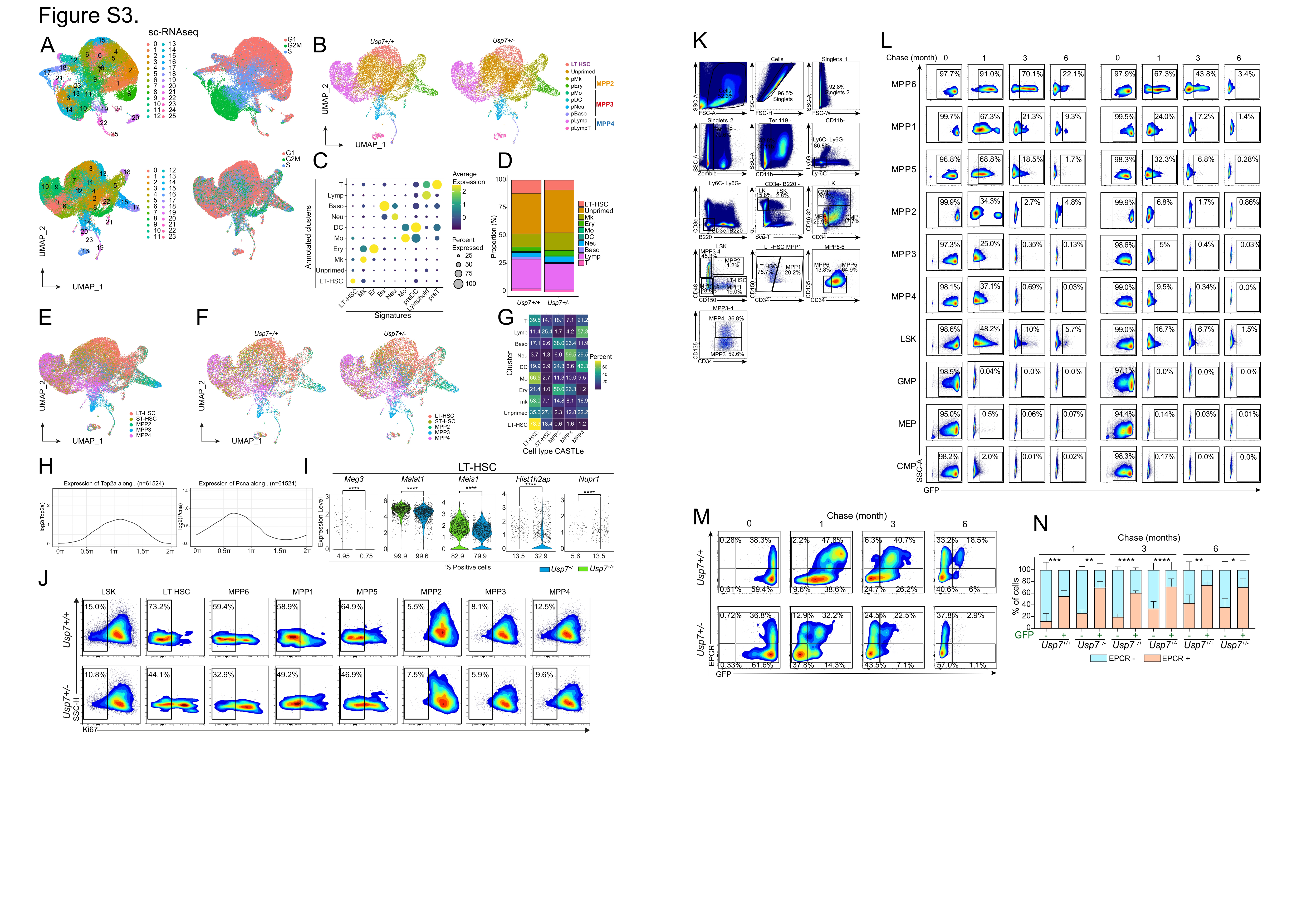

### Supplemental Fig S4

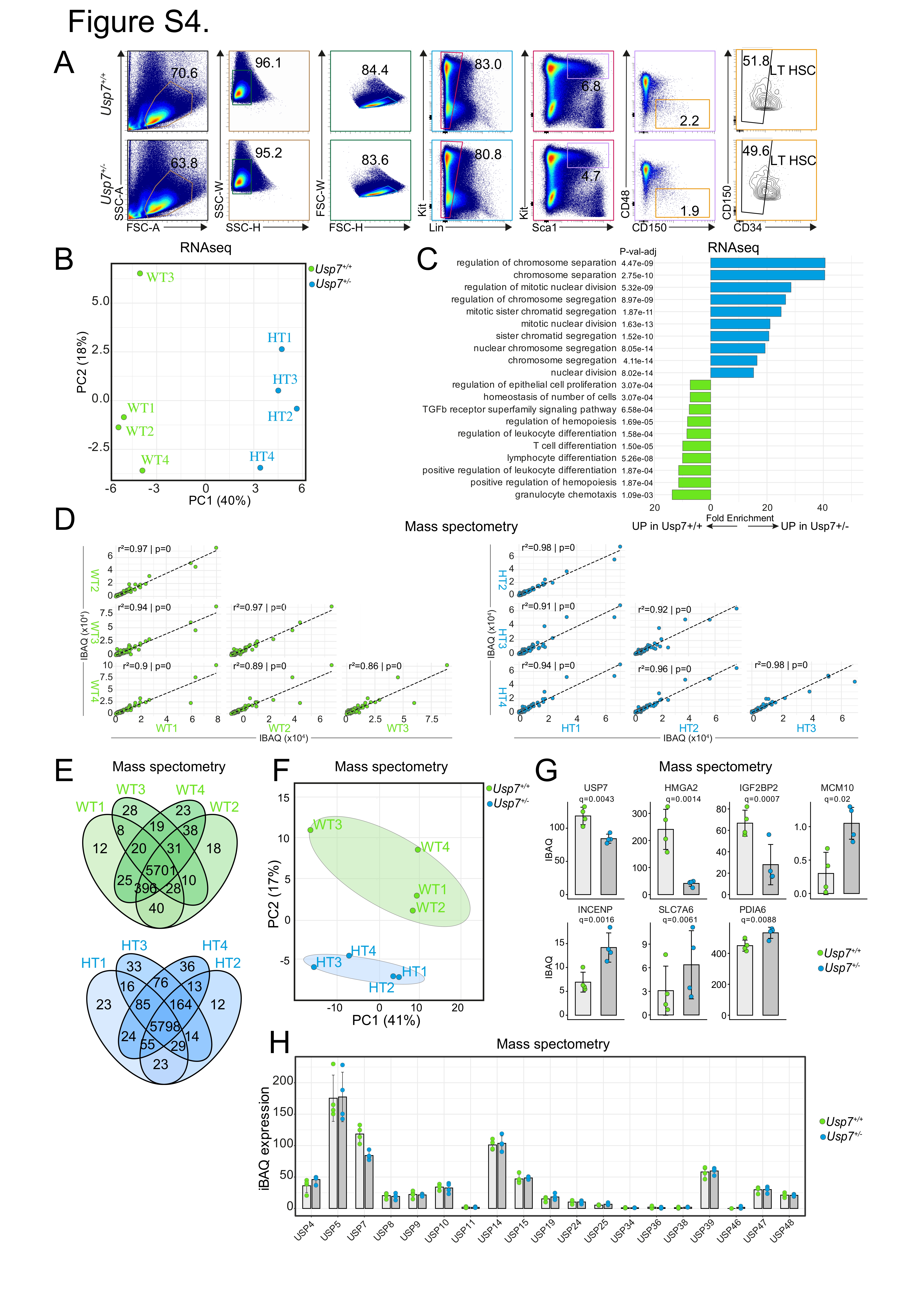

### Supplemental Fig S5

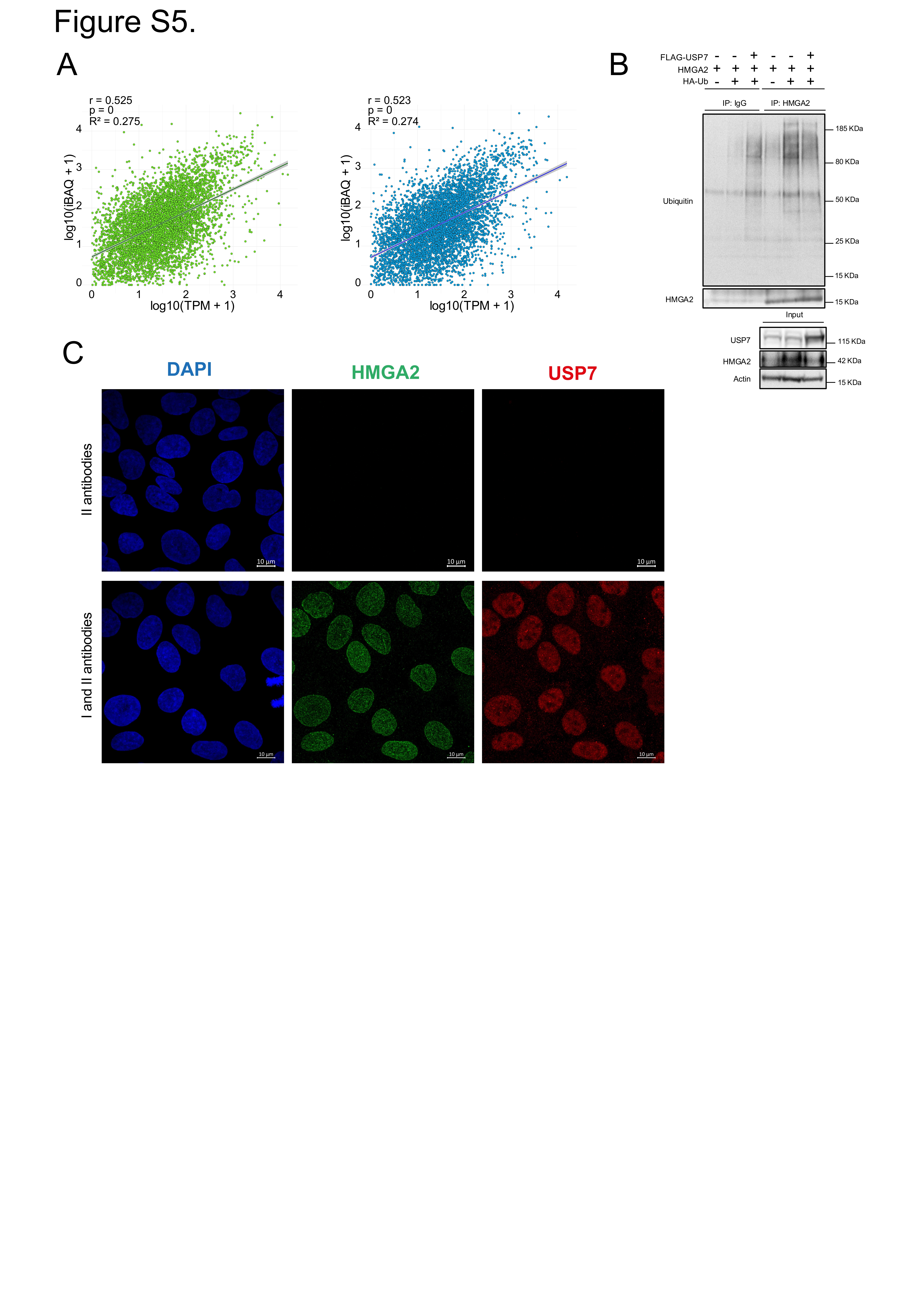
